# Fast two-photon volumetric imaging of an improved voltage indicator reveals electrical activity in deeply located neurons in the awake brain

**DOI:** 10.1101/445064

**Authors:** Mariya Chavarha, Vincent Villette, Ivan K. Dimov, Lagnajeet Pradhan, Stephen W. Evans, Dongqing Shi, Renzhi Yang, Simon Chamberland, Jonathan Bradley, Benjamin Mathieu, Francois St-Pierre, Mark J. Schnitzer, Guoqiang Bi, Katalin Toth, Jun Ding, Stéphane Dieudonné, Michael Z. Lin

**Affiliations:** Department of Bioengineering, Stanford University, Stanford, California, USA; Department of Neurobiology, Stanford University, Stanford, California, USA; Institut de Biologie de l′École Normale Supérieure (IBENS), École Normale Supérieure, CNRS, INSERM, PSL Research University, Paris, France; CNC Program, Stanford University, Stanford, California, USA; School of Life Sciences, University of Science and Technology of China, Hefei, China; Department of Neurosurgey, Stanford University, Stanford, California, USA; Department of Psychiatry and Neuroscience, Quebec Mental Health Institute, Université Laval, Québec, Canada; Baylor College of Medicine, Houston, Texas, USA; CAS Center for Excellence in Brain Science and Intelligence Technology, Shanghai, China

**Author notes:** These authors contributed equally to this work (Mariya Chavarha, Vincent Villette, Ivan K. Dimov).

## Abstract

Imaging of transmembrane voltage deep in brain tissue with cellular resolution has the potential to reveal information processing by neuronal circuits in living animals with minimal perturbation. Multi-photon voltage imaging *in vivo*, however, is currently limited by speed and sensitivity of both indicators and imaging methods. Here, we report the engineering of an improved genetically encoded voltage indicator, ASAP3, which exhibits up to 51% fluorescence responses in the physiological voltage range, sub-millisecond activation kinetics, and full responsivity under two-photon illumination. We also introduce an ultrafast local volume excitation (ULOVE) two-photon scanning method to sample ASAP3 signals in awake mice at kilohertz rates with increased stability and sensitivity. ASAP3 and ULOVE allowed continuous single-trial tracking of spikes and subthreshold events for minutes in deep locations, with subcellular resolution, and with repeated sampling over multiple days. By imaging voltage in visual cortex neurons, we found evidence for cell type-dependent subthreshold modulation by locomotion. Thus, ASAP3 and ULOVE enable continuous high-speed high-resolution imaging of electrical activity in deeply located genetically defined neurons during awake behavior.

## INTRODUCTION

The ability to record electrical activity in individual genetically labeled neurons within living animals would greatly facilitate efforts to understand how nervous systems represent and process information^1^. Fluorescent genetically encoded voltage indicators (GEVIs) can reveal non-spiking electrical activity and resolve action potential (AP) timing with sub-millisecond resolution, tasks that cannot be performed by fluorescent genetically encoded calcium indicators (GECIs)^2^. Multi-photon microscopy, by selectively exciting fluorescence only at the focal point, suppresses the generation of background fluorescence and enables segregation of signals between cells at deeper locations than one-photon microscopy, and is essential for imaging GECIs in many regions of the brain^3^. Despite the unique capabilities of GEVIs and multi-photon microscopy, however, they have yet to be used together for single-trial detection of single APs or subthreshold activity in individual neurons within the live mammalian brain.

Indeed, fast multi-photon imaging of voltage has been considered exceedingly difficult or impractical based on calculations of multi-photon imaging framerates^4,5^ and the limited number of GEVI molecules in one optical section^4,6^. Due to the focal nature of multi-photon excitation, multi-photon imaging is typically performed by raster scanning over cells of interest, yielding sampling rates of ≤ 30 Hz. While this approach is sufficient to capture GECI responses, which decay over hundreds of milliseconds, GEVIs require sampling rates 1–2 orders of magnitude faster if precise tracking of APs or fast oscillations is desired. Multi-photon optogenetic stimulation encounters a similar need for speed, which has been addressed by parallel excitation strategies involving holographic focal volume shaping^7,8^. However, parallel excitation is problematic for imaging, as scattered fluorescence signals from simultaneously excited cells would become intermixed, hindering detection of small responses. Furthermore, as GEVIs reside in the membrane rather than the cytosol, the number of indicator molecules that can be excited in an optical section through a mammalian cell body is typically smaller for GEVIs than for GECIs^4,6^.

Given these limitations, the development of GEVIs with larger two-photon responses to electrical events of interest is highly desirable. Currently, the GEVIs with the largest responses to both subthreshold changes and APs are based on two types of voltage-sensing domains: seven-transmembrane helix opsins and four-transemembrane helix voltage-sensing domains (VSDs). Opsin-based GEVIs have been used *in vivo* with one-photon excitation to report electrical activity of superficially located neurons^9,10^, but their responsivity is severely attenuated under two-photon excitation^4,11^. In contrast, ASAP-family GEVIs, composed of a circularly permuted green fluorescent protein variant inserted within the VSD of *G. gallus* voltage-sensing phosphatase (**Fig. 1a**), are fully responsive under two-photon excitation^11^. In particular, ASAP2s demonstrates the largest response per AP of fluorescent protein-based GEVIs, but its kinetics are actually slower than earlier ASAP variants^11^. If ASAP-family kinetics and/or overall responsivity could be improved, then electrical events could be more easily detected by two-photon imaging.

**Figure 1.**
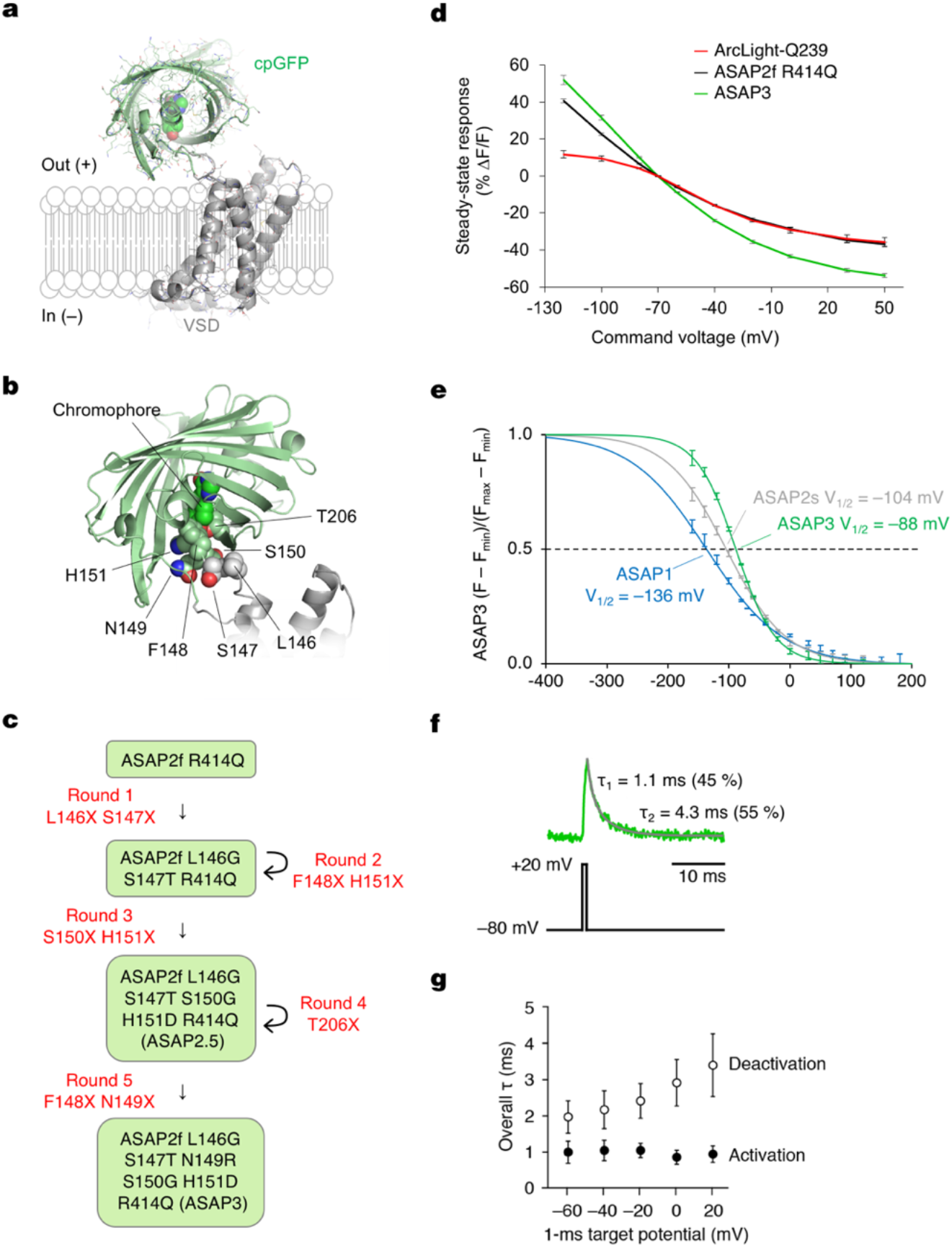
Directed evolution of improved ASAP indicators. **a**, Model showing ASAP domain organization. **b**, The junction site between S3 and cpGFP, with residues targeted for mutagenesis labeled. **c**, Summary of the screening rounds to generate ASAP3. **d,** Fluorescence responses of ArcLight-Q239 (n = 3), ASAP2f R414Q (n = 6), and ASAP3 (n = 10) to 500-ms voltage steps from –120 to 50 mV were characterized by simultaneous patch-clamp and optical imaging at 200 Hz. **e,** Characterization of fluorescence-voltage slope relationship (F-V curves) for ASAP variants. Normalized sigmoid traces fit to steady-state fluorescence responses of ASAP1 (blue trace, n = 6 and 4), ASAP2s (grey, n = 5 and 7), or ASAP3 (green, n = 3 and 6) to 500-ms depolarizing (–70 to +180 mV) and hyperpolarizing (0 to –180 mV) voltage steps. Error bars are SEM. **f,** Example ASAP3 fluorescence trace during 1-ms steps between –80 and +20 mV at 33 °C. Activation in response to the 1-ms step occurs with a peak < 1 ms later. **g,** ASAP3 activation in response to 1-ms pulses at 33 °C occurs with submillisecond kinetics, while deactivation is slower. For deactivation from each voltage level, a single weighted time constant *τ* was calculated as (*a_1_τ_1_* + *a_2_τ_2_*) / (*a_1_ + a_2_*), where *a_1_* and *a_2_* are the coefficients for the biexponential fit with time constants *τ1* and *τ2.* Circles and error bars represent mean ± SD for n = 16 cells. Normalized F–V curves highlight steepening and right-shifting of ASAP3 curve, with both effects contributing to its improved dynamic range. Error bars are SEM.

Here we report an improved indicator, ASAP3, resulting from novel methods for generation and screening of GEVI libraries in mammalian cells. ASAP3 features the largest responses of fluorescent GEVIs to either steady-state voltages or APs under either one- or two-photon excitation, favorable kinetics for both AP timing and detection, and efficient membrane localization. We also introduce an ultrafast local volume excitation (ULOVE) strategy for two-photon microscopy to selectively excite large membrane areas of ASAP3-expressing cells at sampling rates of up to 15 kHz per cell. Using ASAP3 and ULOVE two-photon microscopy in awake head-fixed mice, we demonstrate optical voltage recordings in deeply located cortical and hippocampal pyramidal neurons. We demonstrate the ability to detect APs and subtreshold voltage changes in single trials, with sub-millisecond temporal resolution, over durations of minutes, and across multiple days.

## RESULTS

### A novel electroporation-based screen for improved GEVIs

High-throughput screening of GEVIs has been challenging due to a lack of methods for reproducibly and quickly inducing membrane potential changes while imaging. Patch-clamping, the standard method for controlling membrane potential with fast kinetics, is laborious and low-throughput. Other methods for modulating voltage have not been able to discriminate between fast and slow GEVIs (**Supplementary Note 1**). We hypothesized that electroporation could be used to induce near-instantaneous membrane depolarization. We built a robotic system to deliver electrical pulses to HEK293-Kir2.1 cells, which have a neuron-like resting membrane potential of –75 mV^12^, during high-speed imaging in multi-well plates (**Supplementary Fig. 1a**), then optimized pulse parameters to achieve reliable electroporation (**Supplementary Fig. 1b-d, Supplementary Note 1**). Responses of ASAP1 and ArcLight to electroporation **Supplementary Fig. 1e)** matched their responses to depolarizing steps from –70 to 0 mV imposed by a patch-clamp electrode (**Table 1)**, and ASAP1 response reached steady state more quickly, consistent with its faster kinetics **(Supplementary Table 1**).

**Table 1.** Characteristics of GEVIs used for single-trial imaging in mammalian brain.

|  | ArcLight Q239<sup>a</sup> | Ace2N-4aa-mNeon<sup>b</sup> | ASAP2s<sup>c</sup> | ASAP3<sup>c</sup> |
| --- | --- | --- | --- | --- |
| Activation kinetics at 22–23 °C |  |  |  |  |
| Fast time constant ($\tau_{\text{fast}}$) | 28 ms | 0.37 ± 0.08 ms | 7.0 ± 0.1 ms | 3.7 ± 0.1 ms |
| Fast component amplitude | 39% | 58 ± 5% | 77 ± 3% | 81 ± 1% |
| Slow time constant ($\tau_{\text{slow}}$) | 271 ms | 5.5 ± 1.4 ms | 79 ± 4 ms | 48 ± 4 ms |
| Deactivation kinetics at 22–23 °C |  |  |  |  |
| Fast time constant ($\tau_{\text{fast}}$) | 104 ms | 0.50 ± 0.09 ms | 16.7 ± 0.3 ms | 16.0 ± 0.3 ms |
| Fast component amplitude | 61% | 60 ± 6% | 65 ± 4% | 81 ± 1% |
| Slow time constant ($\tau_{\text{slow}}$) | 283 ms | 5.9 ± 0.9 ms | 116 ± 11 ms | 102 ± 2 ms |
| Activation kinetics at 33–35 °C |  |  |  |  |
| Fast time constant ($\tau_{\text{fast}}$) | 9 ± 1 ms | ND | 1.50 ± 0.13 ms | 0.94 ± 0.06 ms |
| Fast component amplitude | 50 ± 3% | ND | 67 ± 2% | 72 ± 1% |
| Slow time constant ($\tau_{\text{slow}}$) | 48 ± 4 ms | ND | 7.80 ± 0.41 ms | 7.24 ± 0.38 ms |
| Deactivation kinetics at 33–35 °C |  |  |  |  |
| Fast time constant ($\tau_{\text{fast}}$) | 17 ± 1 ms | ND | 3.40 ± 0.37 ms | 3.79 ± 0.15 ms |
| Fast component amplitude | 79 ± 3% | ND | 63 ± 9% | 76 ± 5% |
| Slow time constant ($\tau_{\text{slow}}$) | 60 ± 7 ms | ND | 13.10 ± 2.11 ms | 16.00 ± 1.21 ms |
| Steady-state $\Delta F/F$ per 100 mV at 22–23 °C | ~35% | ~-9% | -36 ± 2% | -51 ± 1% |
| Single-AP $\Delta F/F$ at 22–23 °C | -3.2 ± 2.2% | ~-5% | -7 ± 1%<br>-12 ± 1%<sup>d</sup> | -18 ± 1% |

GEVIs that have been used to record membrane voltage in single trials in live mammals or acute mammalian brain slice are included for comparison. Note only population measurements have been reported for ArcLight Q239 in mammals. For standardized comparisons across studies, kinetics measured in HEK293 or CHO cells in response to step voltages from a -80 or -70 mV baseline to +20 or +30 mV, and AP responses measured in cultured rat hippocampal neurons with a sampling rate of 1–2 kHz, are listed. Responses had reached steady state before voltage was returned to baseline for measurement of deactivation kinetics. Values are mean ± SEM. <sup>a</sup>From ref. 42, with n = 6 cells for kinetics and 20 neurons for APs, except for kinetic parameters at 22–23 °C, which are from ref. 43. <sup>b</sup>From ref. 9, n = 6 cells for kinetics. <sup>c</sup>From this study unless otherwise noted, with n = 9 or 10 cells for kinetics and steady-state responses at 22 °C, n = 7 or 16 cells for kinetics and steady-state responses of ASAP2s or ASAP3 respectively at 33 °C, and n = 4 or 5 neurons for APs. <sup>d</sup>From ref. 11, with n = 5 neurons. ND, not determined.

As retrieving stochastically introduced DNA from single identified cells after electroporation would be difficult, while testing each mutant across many cells would improve statistical precision, we chose to conduct screening in multi-well plates. For this, we devised a fast protocol for multi-well library construction and screening with several unique features. First, we generated complete mutant genes by PCR and transfected the PCR products into cells, with all steps in 384-well format. Previously described GEVI screens utilized plasmid transfection, but this introduces expensive and time-consuming steps of plasmid assembly, transformation, culture, and purification^13–17^. We found that transfection of PCR-assembled genes (**Supplementary Fig. 2a,b**) produced per-cell expression levels similar to plasmids (**Supplementary Fig. 2c**), and that purification of the PCR products was not necessary (**Supplementary Fig. 2d**). Second, unlike previous GEVI screens^13–17^, we produced all mutants deterministically, e.g. performing 400 PCR reactions for all combinations at two sites. This eliminated the need for oversampling to account for Poisson sampling statistics, thereby assuring complete coverage and improving efficiency. Finally, we devised MATLAB routines to perform electroporation, image capture, and image analysis (**Supplementary Note 2, Supplementary Fig. 2e-g**). The automated analysis successfully distinguished sensors with distinct responsivity and kinetics (**Supplementary Fig. 2h-j**). The combination of direct transfection of PCR products and automated screening and analysis enables the evaluation of thousands of mutants daily. This represents an improvement in screening rate of two orders of magnitude compared to patch-clamp electrophysiology.

### Mechanism-based evolution of ASAP3

To generate an optimal ASAP-family template for library-based screening, we first explored combining known beneficial mutations. The response of a GEVI to various electrical waveforms depends on both its steady-state fluorescence-voltage relationship and its response kinetics^2^. Larger response amplitudes and faster activation kinetics are advantageous, as they improve the detection and timing of both APs (which have durations of < 5 ms) and slower events such as excitatory post-synaptic potentials (EPSPs)^18^. ASAP2f differs from ASAP1 by replacement of two amino acids in the linker between the S3 helix of the VSD and the cpGFP domain (pictured in **Fig. 1a**) with one optimized amino acid (aa), resulting in larger responses to hyperpolarization^19^. In ASAP2s, a mutation of Arg-415 to Gln (R415Q, numbering based on contiguity with *G. gallus* voltage-sensing phosphatase for consistency with ASAP1) improved steady-state responsiveness to depolarization by 66%, but reduced its speed (**Table 1, Supplementary Table 1**)^11^. The combined mutant, ASAP2f R414Q (in which aa 414 corresponds to aa 415 in ASAP2s), exhibited responsivity similar to ASAP2s but slightly slower kinetics (**Supplementary Fig. 3a,b, Supplementary Table 1**). However, we chose it as the mutagenesis template because its shorter S3-cpGFP linker reduces the sequence search space as well as the physical degrees of freedom between VSD and cpGFP domains, which should be favorable for enhancing the conformational coupling between these domains.

To identify high-yield sites in ASAP2f R414Q for mutagenesis, we postulated that conformational changes in the voltage-sensing domain affect the pKa of the cpGFP chromophore, similar to the mechanisms of calcium sensing by GCaMP GECIs^20^. Specifically, voltage-induced conformational changes in the VSD may be transduced from the S3-cpGFP linker through the first three positions of cpGFP to influence the position of His-151 (His-148 in wild-type GFP), which normally stabilizes the deprotonated (blue-absorbing) state of the chromophore via hydrogen bonding. We thus decided to target aa 146–151, beginning in the S3-cpGFP linker and ending at His-151 (**Fig. 1b**), choosing one or two sites for saturation mutagenesis in each round.

Over five rounds of screening, four of which involved combinatorial mutagenesis at two positions (**Fig. 1c, Supplementary Figs. 3-7, Supplementary Note 2**), we obtained a variant that responded to depolarization from – 70 to +30 mV with a fluorescence change (*∆F/F*) of –50.9 ± 1.1% (mean ± standard error of the mean (SEM), n = 10 cells). This is the largest fluorescence response among fluorescent protein-based voltage indicators characterized so far, representing a 46% improvement over the –34.9 ± 1% (n = 6 cells) of ASAP2f R414Q (**Fig. 1d**). We named this variant, which differs from ASAP2f by the mutations L146G S147T N149R S150G H151D R414Q, as ASAP3.

### Detailed characterization of ASAP3

To understand the basis for the improved responsiveness of ASAP3, we first compared its input-output function, i.e. its fluorescence–voltage (F–V) curve, to that of earlier variants. ASAP1, ASAP2s, and ASAP3 responses were well fit by sigmoidal Boltzmann functions, similar to the responses of other VSD-based GEVIs and to the gating charge movements of the VSDs themselves^21^ (**Supplementary Fig. 8a**). Interestingly, ASAP3 demonstrates a larger maximal fluorescence change across its extrapolated voltage sensitivity range (4.5-fold) than ASAP2s or ASAP1 (3.3-fold each), due to more complete dimming at more positive membrane voltages (**Supplementary Fig. 8a**). Normalized F–V curves revealed progressive stabilization of the bright state from ASAP1 to ASAP2s and finally to ASAP3. This produced a narrowing of the voltage tuning range and a progressive right-shifting of the midpoint response value (V1/2), from –136 mV in ASAP1 to –104 mV in ASAP2s and –88 mV in ASAP3 (**Fig. 1e**). As these V1/2 values all reside to the left of the physiological voltage range of –70 to +30 mV, narrowing and right-shifting the F-V curve has the desirable effect of steepening responses to physiological events.

We next characterized kinetics of ASAP3 responses. To improve the reliability of spike detection, it is favorable to have activation kinetics that are as fast as possible and deactivation kinetics that are delayed, which maximizes the integrated fluorescence change to a single spike^23^. To allow comparisons to previous measurements of other GEVIs, we first tested complete responses (reaching steady state) to voltage steps from –70 to +30 mV at 22 °C. The time course of ASAP3 activation was well fit to a biexponential curve (**Supplementary Fig. 8b**), with a fast time constant of 3.7 ± 0.1 ms (mean ± SEM, n = 12 cells), significantly shorter than the 7.0 ± 0.1 ms measured for ASAP2s (**Table 1**). The fast component accounted for 81 ± 1% of the activation response for ASAP3, similar to the 77 ± 3% of ASAP2s (**Table 1**). Kinetics of ASAP3 were also faster than ASAP2s in response to hyperpolarizing steps (**Supplementary Table 1**). To understand ASAP3 and ASAP2s performance under more physiological conditions, we also characterized kinetics at 33 °C. The complete response to a 50-ms depolarizing step at 33 °C exhibited a fast time constant of 0.94 ± 0.06 ms (mean ± SEM, n = 16 cells) with amplitude 72 ± 1%, 1.6-fold faster than ASAP2s (1.50 ± 0.13 ms, amplitude 67 ± 2%; n = 7 cells) (**Supplementary Fig. 8c, Table 1**). Finally, we characterized the kinetics of responses to 1-ms voltage steps from –80 to +20 mV at 33 °C to mimic APs (**Supplementary Table 2**). Consistent with its faster kinetics, ASAP3 reached 54 ± 1% (mean ± SEM, n = 16 cells) of its total responsivity during the 1-ms step, significantly higher than ASAP2s at 39 ± 1% (mean ± SEM, n = 7 cells). Thus, ASAP-family kinetics are approximately 4-fold faster at 33 °C than at 22 °C, with ASAP3 exhibiting a sub-millisecond activation time constant.

Interestingly, ASAP3 preserved the relatively slow return to baseline of ASAP2s, which is useful for prolonging the response to an AP and thereby facilitating the identification of spikes by template matching^11^. At 33 °C, ASAP2s fluorescence returned to baseline following a 1-ms step in a biexponential manner, with time constants of 2.11 ± 0.40 ms (amplitude 53 ± 10%) and 7.10 ± 1.62 ms (**Supplementary Table 2**). Deactivation kinetics for ASAP3 were similar with time constants of 1.48 ± 0.09 ms (amplitude 60 ± 3%) and 6.30 ± 0.40 ms (**Fig. 1f**), resulting in an overall weighted deactivation time constant of 3.41 ± 0.21 ms (**Fig. 1g**). Deactivation kinetics for both ASAP2s and ASAP3 were slower following prolonged depolarization compared to the 1-ms step, and were further slowed by reducing the temperature to 22 °C (**Table 1**).

The 46% larger responsivity and 60% faster activation kinetics of ASAP3 compared to ASAP2s should improve spike detection in neurons in a multiplicative manner, as long as expression levels and membrane localization are not impaired. In cultured hippocampal neurons, ASAP3 efficiently localized to the plasma membrane (**Fig. 2**), as seen with previous ASAP variants^11,18,19^. ASAP3 fluorescence outlined dendritic spines, and all ASAP3 signal throughout the neuron responded to voltage steps (**Fig. 2a, Supplementary Movie 1**). Brightness of ASAP3 at rest was comparable to ASAP2s (**Supplementary Fig. 9**). Responses to current-evoked APs (**Fig. 2b**) were indeed larger with ASAP3, reaching 2.7-fold higher peak amplitude than ASAP2s (*∆F/F* of −17.9 ± 1.2% vs −6.6 ± 0.8% respectively) and 2.4-fold larger integrated response (**Fig. 2c**). Extended recordings in neurons confirmed the ability of ASAP3 to track both subthreshold depolarizations and APs, with > 20% responses to APs in some cases (**Fig. 2d**). Thus, the reporting of APs in neurons is dramatically improved with ASAP3 compard to ASAP2s.

**Figure 2.**
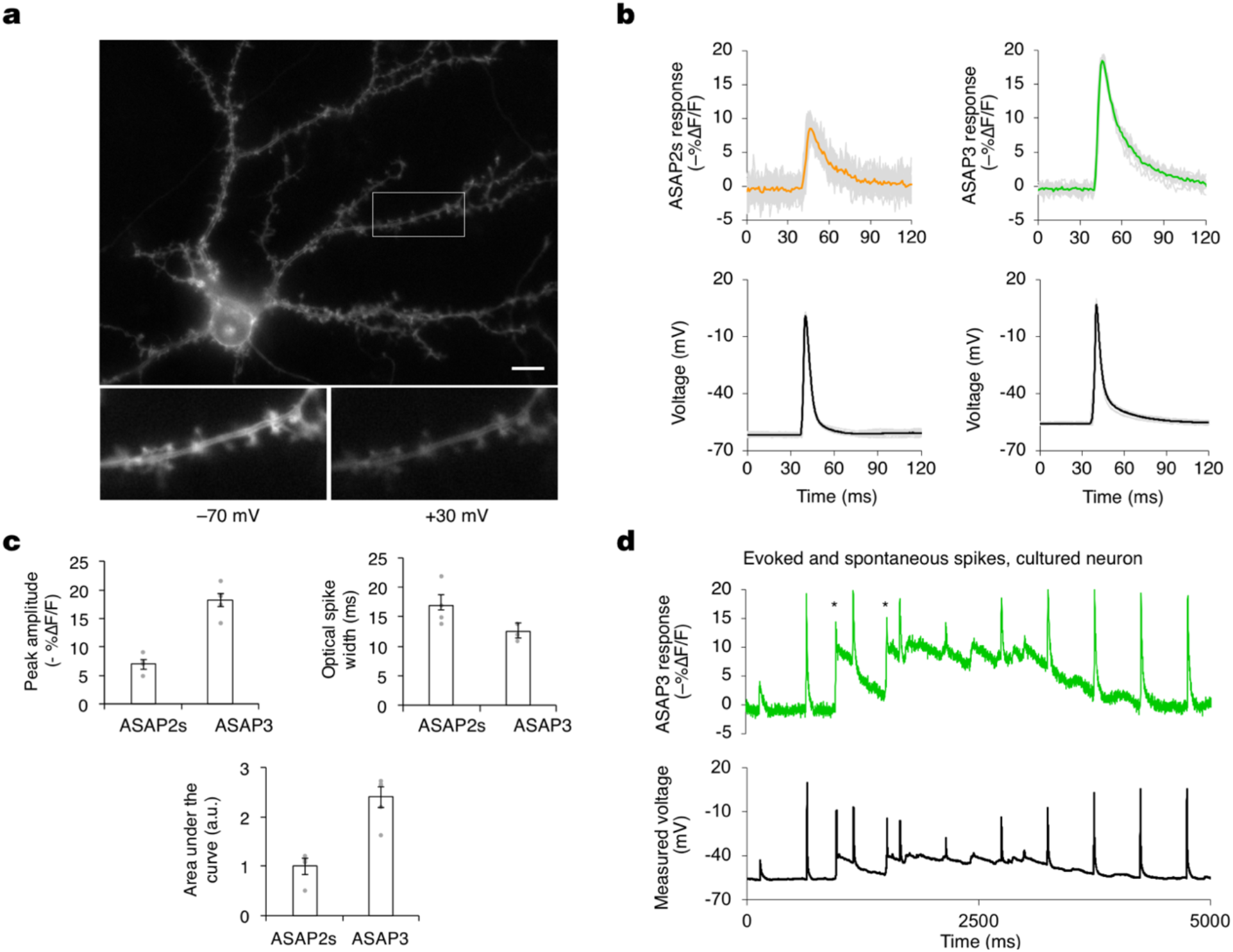
Characterization of ASAP3 in neurons. **a**, ASAP3 is effectively targeted to the plasma membrane in a representative neuron imaged during voltage-clamp electrophysiology. Higher magnification views of the boxed region in panels below show voltage-dependent decrease in fluorescence from a dendritic region including spines. Scale bar is 10 μm. **b,** Responses of ASAP2s and ASAP3 to current-evoked action potentials in representative neurons selected for their matched AP waveforms: peak amplitudes of 63.5 ± 0.8 mV and 63.7 ± 1.0 mV; full width at half maximum (FWHM) of 5.0 ± 0.1 and 3.9 ± 0.2 ms for ASAP2s and ASAP3 respectively. Single trial responses (grey, n = 25 for each construct) were averaged to give mean fluorescence responses. **c,** Mean peak fluorescence responses, optical spike width at half-maximal height, and areas under the curve of fluorescence responses were determined for ASAP2s (n = 4 cells) and ASAP3 (n = 5 cells). Three to 34 APs were recorded per neuron, and were averaged to determine the contribution from each cell. APs for ASAP2s and ASAP3 reached 10.2 ± 3.3 mV and 8.5 ± 2.1 mV peak voltages respectively, and had the same width of 4.4 ± 0.3 ms in the voltage trace at half-maximal height. Areas under the curve were determined by integrating fluorescence response curves from 40 to 80 ms. Values are mean ± SEM. **d**, Electrical and optical responses from a representative neuron expressing ASAP3 to current-triggered APs. Asterisks indicate spontaneous spikes not elicited by current injection.

### Two-photon imaging of spikes with ASAP3

To characterize ASAP3 responses in neuronal tissue and under two-photon excitation, we patch-clamped neurons in hippocampal slice cultures while exciting ASAP3 with random-access multiphoton (RAMP) microscopy (**Fig. 3a)**. ASAP3 activation kept pace with the rising phase of APs at either 20 °C or ~32 °C, while return to baseline was more prolonged than APs in a temperature-dependent manner (**Fig. 3b**), as expected from the known deactivation kinetics (**Table 1**). ASAP3 produced responses of 17.0 ± 0.8% to single APs at ~32 °C, significantly larger than ASAP2s at 11.7 ± 0.7% (each mean ± SEM, n = 12 neurons), and similar to 1-photon results in cultured hippocampal neurons (**Fig. 3c,d)**. The discriminability index (*d’*), which enables a prediction of false positive and false negative rates by signal detection theory^22^, was 22.5 ± 4.4 (mean ± SEM, n = 3 neurons) for ASAP3 and 12.2 1.32 (n = 11 neurons) for ASAP2s for single APs recorded at 20 voxels per neuron. At the sampling frequency of 925 Hz, a *d′* value of 22.5 implies a theoretical false positive rate of once per 2.6 × 10^22^ hours and a theoretical false negative rate of 1.2 × 10^−29^ per event. These results indicate that ASAP3 can reliably detect spikes with two-photon imaging of a few membrane locations, and is superior to ASAP2s.

**Figure 3.**
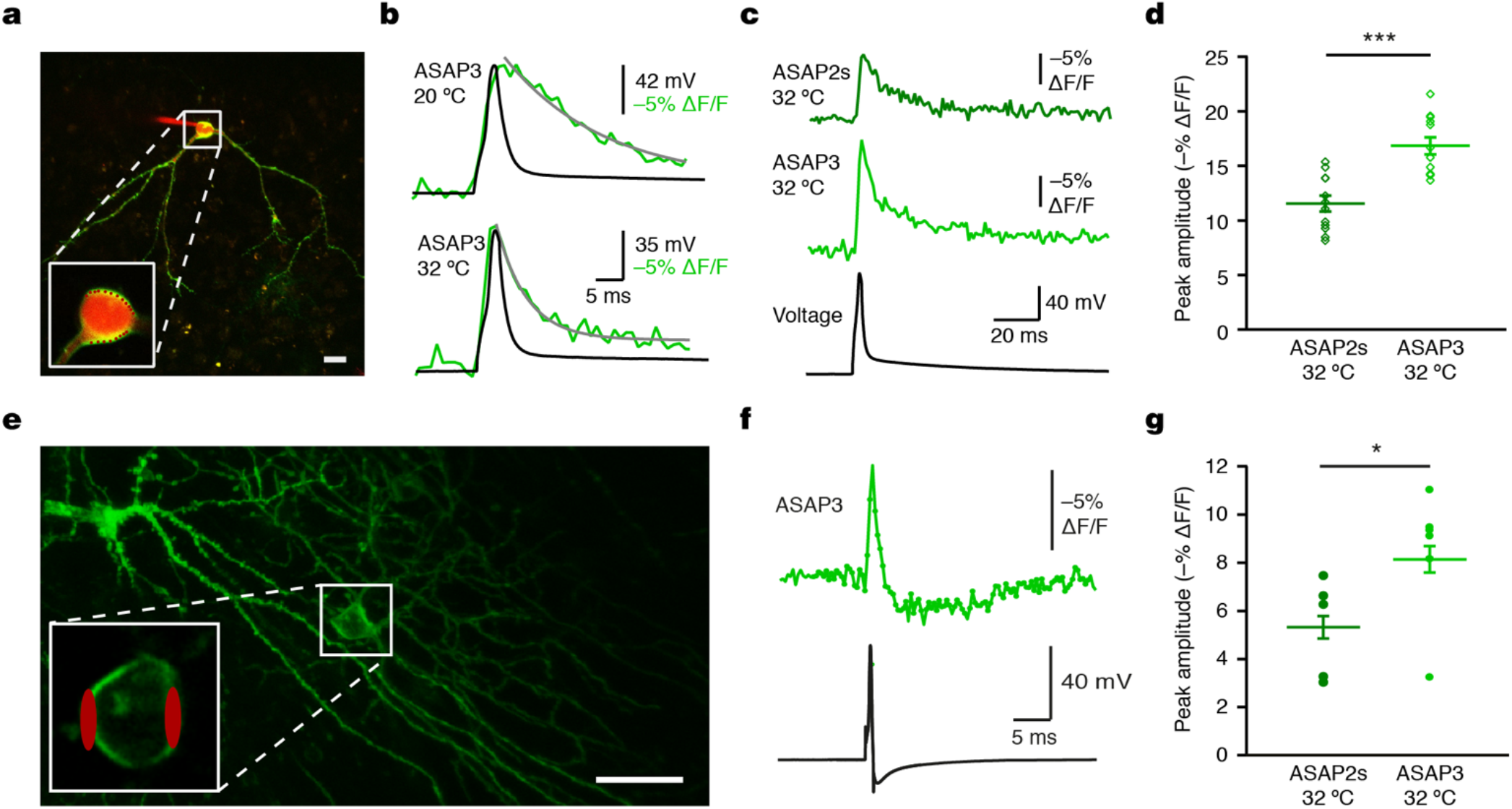
Spike detection in mammalian brain tissue by ASAP3 and two-photon microscopy. **a,** An overlay image of a two-photon maximum-intensity z projection of an ASAP3-expressing neuron recorded in the whole-cell configuration with an intracellular solution containing red fluorescent Alexa Fluor 594. The inset shows the expanded somatic region of the neuron, with red dots marking the 20 recorded voxels. Scale bar, 20 μm. **b**, Representative examples of ASAP3 responses to current-evoked APs recorded at either 20 or 32 °C. The ASAP3 signal tracks the rising phase of AP, but exhibits a longer decay time than the AP. About half of the decay exhibited a time constant of 4.3 ± 0.4 ms (n = 12 neurons) at 32 °C or 9.3 ± 1.9 ms (n = 8) at 20 °C. A second component of about 30 ms accounted for the other half of the decay, which may relate to the decay of the underlying after-depolarization. **c**, Representative examples of ASAP2s and ASAP3 response to current-evoked APs at 32 °C (average of 20 voxels over 10 trials). **d**, ASAP3 shows higher peak amplitude responses than ASAP2s to current-evoked APs recorded at 32 °C. ***p < 0.0001 (two-tailed t-test). Bars represent mean ± SEM. **e**, Two-photon stack projection of sparse ASAP3 expression in molecular layer interneurons in a parasagittal cerebellar slice. The inset shows the expanded somatic region of the neuron, with red ellipsoids marking the recorded regions. Scale bar, 20 μm. **f**, Top, averaged ASAP3 fluorescence transient (50 trials) for the interneuron in **e** shows a characteristic interneuron waveform with after-hyperpolarization. Bottom: corresponding spike shape recorded in whole-cell current clamp mode. **g,** ASAP3 shows improved responsivity over ASAP2s for APs in molecular layer interneurons. Bars represent mean ± SEM. *p = 0.026 (Wilcoxon rank sum test).

To challenge ASAP3 in a difficult detection task, we recorded from molecular layer interneurons in acute cerebellar slices (**Fig. 3e**). These neurons generate very narrow spikes at 34 °C (full width at half maximum of 0.43 ± 0.06 ms, mean ± standard deviation (SD), **Fig. 3f**). ASAP3 still produced *∆F/F* of 8.3 ± 1.1% (mean ± s.em., n = 6 neurons) for spikes in these extreme conditions, outperforming the 5.3 ± 0.9% of ASAP2s (n = 5 neurons) (**Fig. 3f,g)**. Thus, ASAP3 also outperforms ASAP2s in the detection of fast spikes.

For imaging voltage at neuronal cell bodies in densely labelled tissue, it is useful to remove GEVI signal from neurites, as the mesh-like distribution of neuritic signals is difficult to separate from cell bodies^22^. We had previously concentrated ASAP2s signal in neuronal cell bodies by fusion at its C-terminus to the Kv1.2 proximal retention and clustering (PRC) segment ^23^. We confirmed that appending the same PRC segment to ASAP3 also concentrates it in neuronal cell bodies, and does not interfere with the voltage responsivity of ASAP3 (**Fig. 4a–c**). Testing *in vivo* confirmed that ASAP3 and ASAP3-Kv signals are exclusively membrane-bound, so that all fluorescence should be responsive to voltage changes, with ASAP3-Kv fluorescence enriched in the cell bodies (**Fig. 4d**).

**Figure 4.**
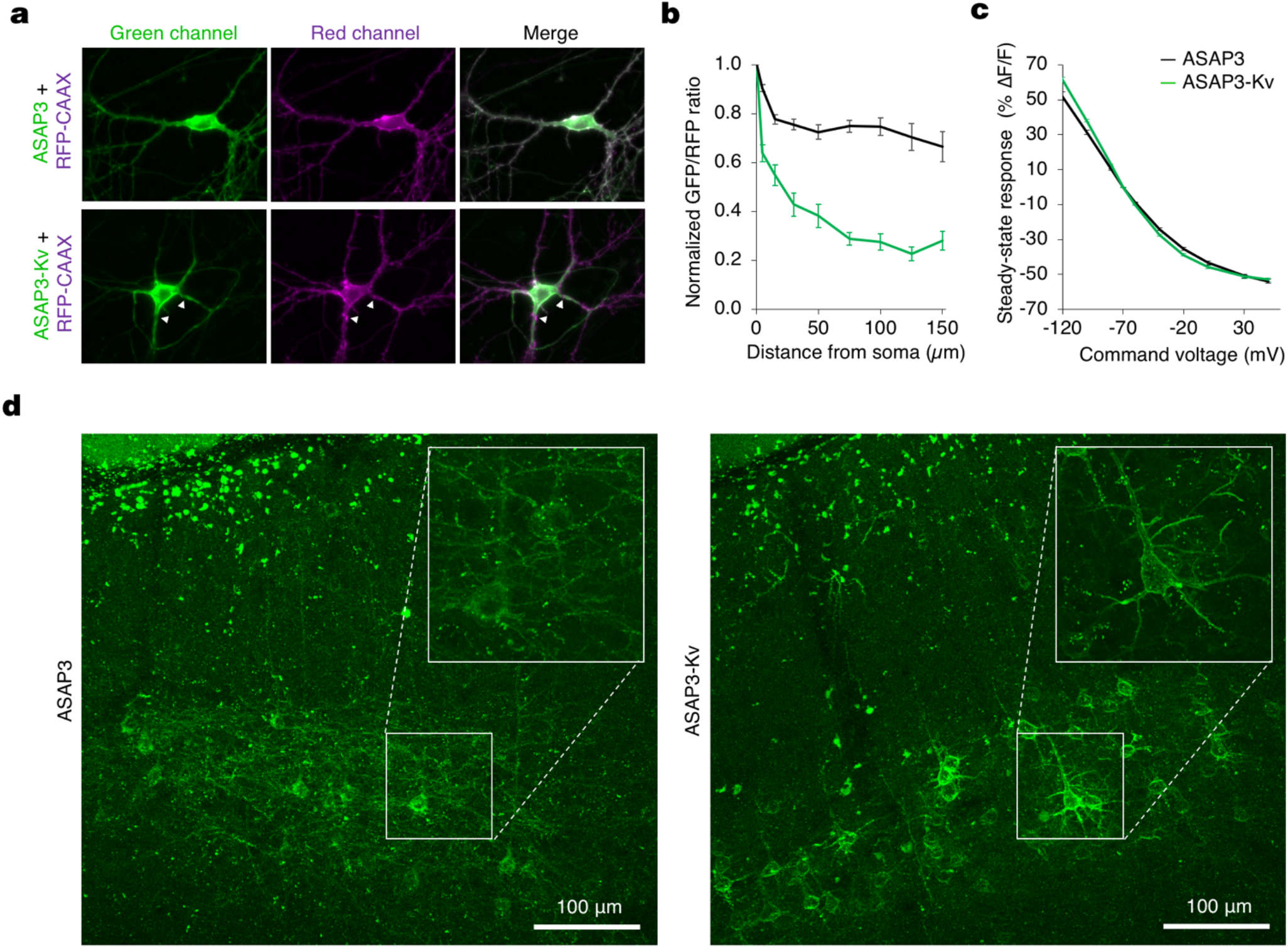
Soma-targeted ASAP3-Kv. **a,** Expression patterns of ASAP3 (green, top) and ASAP3-Kv (green, bottom) compared to an RFP-CAAX membrane marker (magenta). ASAP3-Kv shows reduced expression in distal dendrites. Interestingly, in proximal dendritic segments ASAP3-Kv can be detected along straight regions of the dendritic membrane but is excluded from spines that are visualized with RFP-CAAX (arrows). **b**, The ratio of ASAP3 (n = 26) or ASAP3-Kv (n = 33) fluorescence to RFP-CAAX fluorescence was quantified at various distances from the soma. **c**, F-V curve of ASAP3-Kv (n = 13) compared to ASAP3 (n = 10). **d**, Representative projection confocal images of ASAP3 (left) and ASAP3-Kv (right) expression in cortical slices reveal that the Kv1.2 PRC motif enriches ASAP3 signals in cell soma.

To characterize the performance of ASAP3-Kv in acute brain slices, we expressed ASAP3-Kv in mouse striatum with adeno-associated virus (AAV), then prepared slices for simultaneous patch-clamp electrophysiology and two-photon imaging. With line-scanning along the edge of neuronal cell bodies at 1 kHz, thereby detecting fluorescence at several voxels with each scan, ASAP3-Kv reported single current-evoked APs at 22 °C with *∆F/F* of −37.3 ± 4.1% (mean ± SD, n = 8 spikes) and a signal-to-noise ratio of 2.6 ± 0.39 (**Supplementary Fig. 10**). As expected, the fluorescence transients peaked simultaneously with the APs but were more prolonged in duration, resulting in a *d’* value of 13.5 ± 2.3. At a sampling frequency of 1 kHz, this value suggests a theoretical false-positive rate of once per 37,577 hours while preserving a theoretical false negative rate of 7.4 × 10^−12^ per event. ASAP3 nevertheless deactivated rapidly enough to enable the discrimination of individual spikes in high-frequency spike trains up to 100 Hz (**Supplementary Fig. 10**), a task that cannot be performed by calcium indicators due to the slower kinetics of both calcium and calcium indicators^2^. Thus, ASAP3-Kv demonstrates high reliability for two-photon detection of single voltage spikes and tracking of high-frequency spike trains in mammalian brain tissue.

### Imaging voltage in deep neurons in awake behaving mice with ASAP3-Kv and ULOVE two-photon microscopy

While RAMP microscopy provides the kilohertz sampling rates needed for spike detection, its application to voltage imaging *in vivo* has been hindered by the small number of GEVI molecules present per voxel and by brain movement. To address these issues, we developed a variant of RAMP microscopy in which driving the acousto-optic deflectors (AODs) with a non-stationary frequency scans local volumes of several tens of cubic microns surrounding a target point within tens of microseconds (**Fig. 5a**). The local volume size and shape are tuned to encompass a large fraction of the neuron membrane during brain movement (**Fig. 5b,c**). Size tuning does not impact the dwell-time or frame rate of the recordings, as larger volumes are scanned at higher speed by design. This ultrafast local volume excitation (ULOVE) strategy efficiently mitigated movement artifacts during locomotion (**Fig. 5d**). Applying ULOVE two-photon imaging to neurons expressing ASAP3-Kv in living mouse brain improved photon flux, as expected from exciting a larger membrane area. Photobleaching was modest even during continuous recordings at kHz frame rates, with an initial fluorescence drop of 19.6 ± 4.5% (mean ± SD, n = 23 neurons) in the first 10 s followed by a slow decrease at 0.99 ± 0.75% per min for the following 5 min (**Supplementary Fig. 11**).

**Figure 5.**
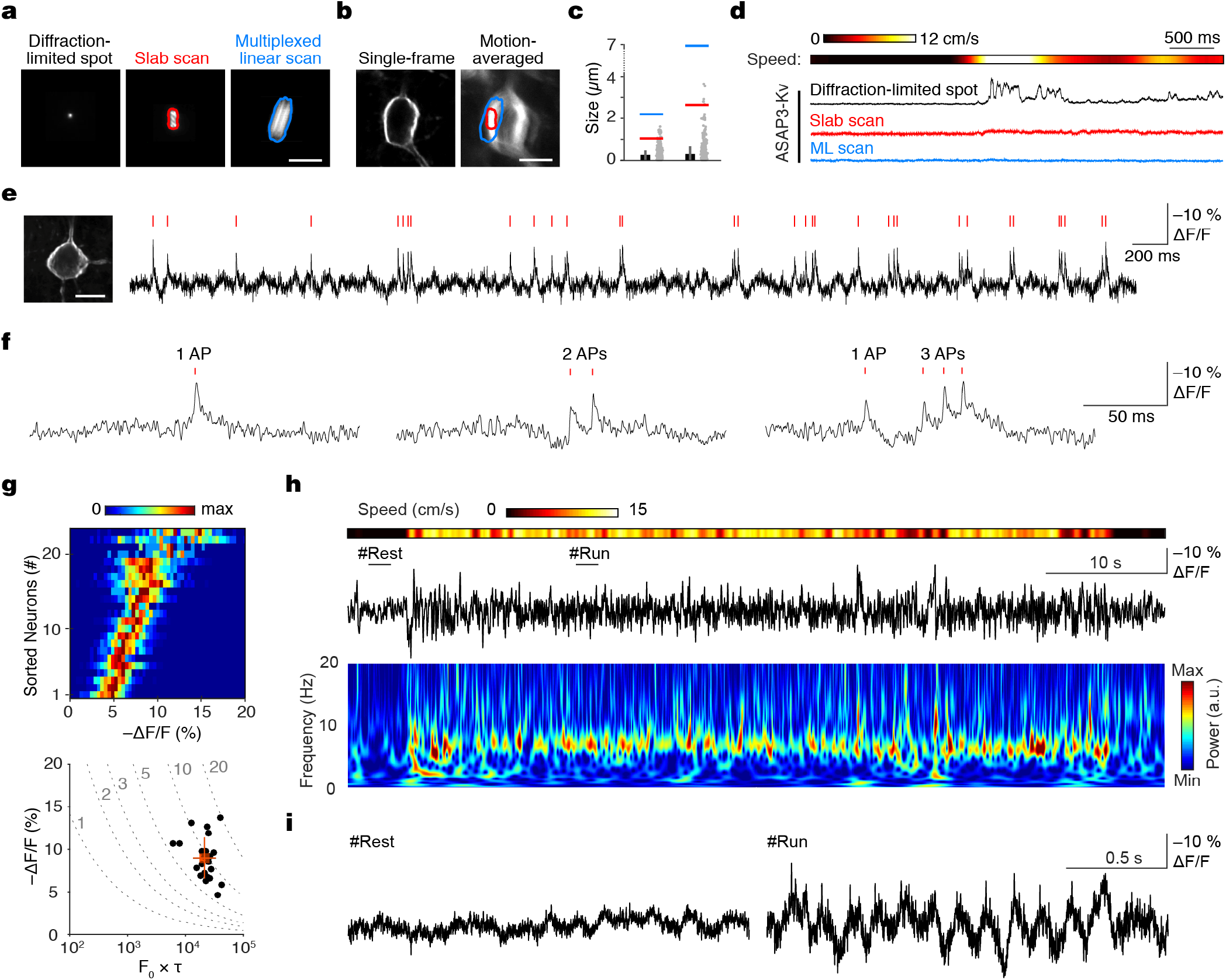
ULOVE two-photon imaging of ASAP3-Kv enables recordings of spikes and subthreshold voltages in awake mice. **a**, Maximal projections of the point spread functions obtained by local volume scanning with AODs and their outlines: left, diffraction limited spot; center, slab scan; right, multiplexed linear scan. Scale bar: 10 μm. **b**, Two-photon timelapse projections of an ASAP3-Kv expressing cortical neuron with (left) and without (right) registration. Volume scanning outlines encompass a large franction of the membrane and movement blur. Scale bar, 10 μm. **c**, Motion-induced displacement calculated from the registration data (grey points) and its mean ± SD (black) (n = 29 recordings). Half-widths of the local volumes are indicated by color-coded lines. **d**, Example of ASAP3-kv signals simultaneously recorded in vivo with the diffraction limited spot (black), slab scan pattern (red) and multiplexed linear scan (blue). Heatmap indicates mouse running speed. **e**, Example recording of ASAP3-Kv-expressing neuron with bursty activity and detected events (red ticks). **f**, High temporal resolution examples of single action potentials and bursts. **g**, Top, heatmap representing the distribution of response amplitudes for detected spikes in the 23 analyzed neurons. Bottom, discriminability index (*d’*) of the 23 neurons displayed on a graph of − *∆F/F* vs. *F_0_∙τ*. Mean and standard deviations are indicated in red. **h,** Fluorescence trace illustrating theta-frequency membrane potential oscillations in a hippocampal neuron (black, top), corresponding time-frequency map (middle), and running speed heat map (bottom). #Rest and #Run, epochs enlarged in b. **i**, Epochs from **h** during rest (top) and run (bottom) behavioral states. Note that fast deflections are nested on hippocampal theta oscillations during run.

We tested the ability of ULOVE two-photon imaging to detect spikes in the visual cortex of awake mice after AAV transduction of ASAP3-Kv. We recorded ASAP3-Kv signals at 3-kHz sampling rates from neurons in layers 1−. To detect spikes, we used the MLSpike algorithm routinely used for GECI signals based on matching to a mono-exponentially decaying template^24^, with 23 cells from five mice passing the statistical criteria for analysis (**see methods)**. Representative ASAP3-Kv signals and spike assignments in a layer-2/3 neuron are shown in **Fig. e,f.** In 150 s of continuous recordings, the sampled neurons overall fired at an average rate of 1.8 ± 2.0 Hz (mean ± SD, range 0.07−10 Hz). ASAP3-Kv transients exhibited *∆F/F* of 9.0 ± 2.4% (mean ± SD, range 4.7–13.6%, **Fig. 5g**). The *d’* value for spikes, calculated from the photon count and ASAP3-Kv waveform of each cell, was 9.3 ± 3.0 (mean ± SD, range 5.9−19.4, **Fig. 5g)**. At a sampling frequency of 3 kHz, a *d′* value of 9.3 ensures a theoretical rate of false positive detection of once every 200 seconds while preserving a theoretical false negative rate of 1.5 × 10^−6^ per event. This provides a large safety margin for *in vivo* experimental conditions, under which subthreshold membrane potential fluctuations and other non-stationary processes arise on top of the photon shot noise.

Unlike calcium indicators, GEVIs can report both subthreshold membrane potential fluctuations, which represent changes in synaptic inputs, and spikes, which represent functional outputs, in the same neuron. Electrical oscillations detected by field recordings or electroencephalography have been recognized as readouts of the synchronized activity of neuronal populations and substrates for neuronal computation^25^, but cannot be detected by calcium imaging. In hippocampal neurons, we observed theta rhythms in the optical ASAP3-Kv signal, which were more frequent during running epochs (peak frequency: 6.95–8.1 Hz, n = 2 cells, **Fig. 5h**) and correlated with running speed (Pearson’s correlation coefficient = 0.25, p < 0.001, **Supplementary Fig. 12a,b**). These fluorescence data closely mimic extracellular local field potential recordings obtained in similar experimental conditions^26^. On top of the voltage oscillation, we detected spikes phase-locked to the depolarizing peak of the theta oscillation (**Fig. 5i, Supplementary Fig. 12c,d**) in line with electrophysiological results. These findings demonstrate the ability of ASAP3 and two-photon imaging to detect subthreshold oscillations and action potentials in the same neuron *in vivo*, a capacity previously restricted to low-yield intracellular and whole-cell patch clamp techniques.

### Realizing unique capabilities of GEVIs with ASAP3-Kv

We took advantage of the ability of two-photon illumination to image voltage in deep locations. Two-photon raster scans through ASAP3-Kv-expressing neurons revealed healthy morphology throughout a cortical column (**Supplementary Movie 2**). We next tested whether voltage dynamics in layer-5 cortical neurons could be detected by ASAP3-Kv and two-photon microscopy. Layer 5 is located ~500 μm below the surface, beyond the 50–100 μm range at which one-photon microscopy can discern individual cells^27,28^. One-photon methods can only image layer-5 cells via inserted optical lenses^29^, which damage adjacent structures and provide s spatially limited coverage of a thin slab of brain tissue. Two-photon microscopy was able to image ASAP3-Kv in layer-5 neurons in visual cortex with high resolution and high contrast without damage to the cortex (**Supplementary Fig. 13a**). ASAP3-Kv signals revealed that 11 out of 12 neurons exhibited membrane potential fluctuations between up and down states, with *∆F/F* of 9.9 ± 2.7% (mean ± SD) between the two states (**Supplementary Fig. 13b)**. Cells spent 12.6 ± 5.7% of their time in up states (**Supplementary Fig. 13c)**, with mean duration of 88 ± 63 ms (n = 1267 up states in 11 cells). These alternations were confirmed by performing two-photon raster scans at the relatively slower framerate of 55 Hz (**Supplementary Movie 3**).

A theoretical advantage of GEVIs is the ability to record voltage in small compartments that are difficult to patch-clamp, such as dendrites. Brightly labeled proximal apical dendrites of layer-5 pyramidal neurons were readily identified in layer 4 (**Supplementary Fig. 13d)**. We reasoned that spikes might be more easily recorded from these dendritic sites, where their duration is increased. Triplet recording revealed uncorrelated spikes in two of three dendrites (**Supplementary Fig. 13e**), while the third displayed up/down states alternations (**Supplementary Fig. 13f)**. Thus volume scanning yields enough signal to perform voltage recordings from dendritic segments, opening the way to the optical study of dendritic integration *in vivo*. Our results also demonstrate that RAMP can image multiple units at high speed in a near-simultaneous interspersed manner.

Finally, GEVIs offer the potential to monitor transmembrane voltage in the same neuron across multiple days (**Supplementary Fig. 14a**), a task that has not been possible to perform with electrodes. Waveforms, detectability and firing properties remained unchanged across day pairs, although individual cells may increase or decrease their firing (**Supplementary Fig. 14b-d**). Thus, we found that ASAP3 and RAMP microscopy achieve multiple theorized advantages of GEVIs, namely the ability to report both subthreshold and spiking electrical activity and the abilities to report from dendritic locations, from multiple units, or over multiple days.

### Modulation of network activity by locomotion in the visual cortex

In the visual cortex, pyramidal cells respond with increased firing rates during locomotion, but the network mechanisms of this effect are not fully understood^30^. Using ASAP3-Kv, we investigated whether locomotion influences synaptic inputs in different neuronal types in visual cortex. ASAP3-Kv spike shape (**Fig. 5e,f, Fig. 6a-d**, **Supplementary Fig. 15a)** and firing behavior of neurons (**Supplementary Fig. 15b**) varied between cells in layers 1–2, as expected from synapsin promoter-mediated expression in multiple cell types. Out of 23 cells, three cells fired only rarely (0.087 ± 0.011 Hz, mean ± SD) and were excluded from further analysis. In the remaining 20 cells, exponential fitting of the interspike interval (ISI) distribution identified 11cells that fired bursts (ISI 23 ± 17 ms, mean ± SD for n = 696 bursts; spikes in bursts 38 ± 20% of total spikes; **Fig. 6e,f**). The spike number distribution in bursts was skewed (**Supplementary Fig. 15c,d**), in agreement with the log-normal distribution expected in cortical networks^31^. All bursty neurons displayed an after-depolarizing potential (*∆F/F* from spike threshold of 5.1 ± 3.2%, mean ± SD, **Fig. 6f**) and fired at low average frequencies (1.8 ± 1.2 Hz), characteristics of pyramidal neurons^32^. In the non-bursty group, six out of nine neurons displayed an after-hyperpolarization (∆F/F of 0.9 ± 0.3% peaking 22.4 ± 6.8 ms after onset, **Fig. 6d,f**), a characteristic of interneurons^32^.

**Figure 6.**
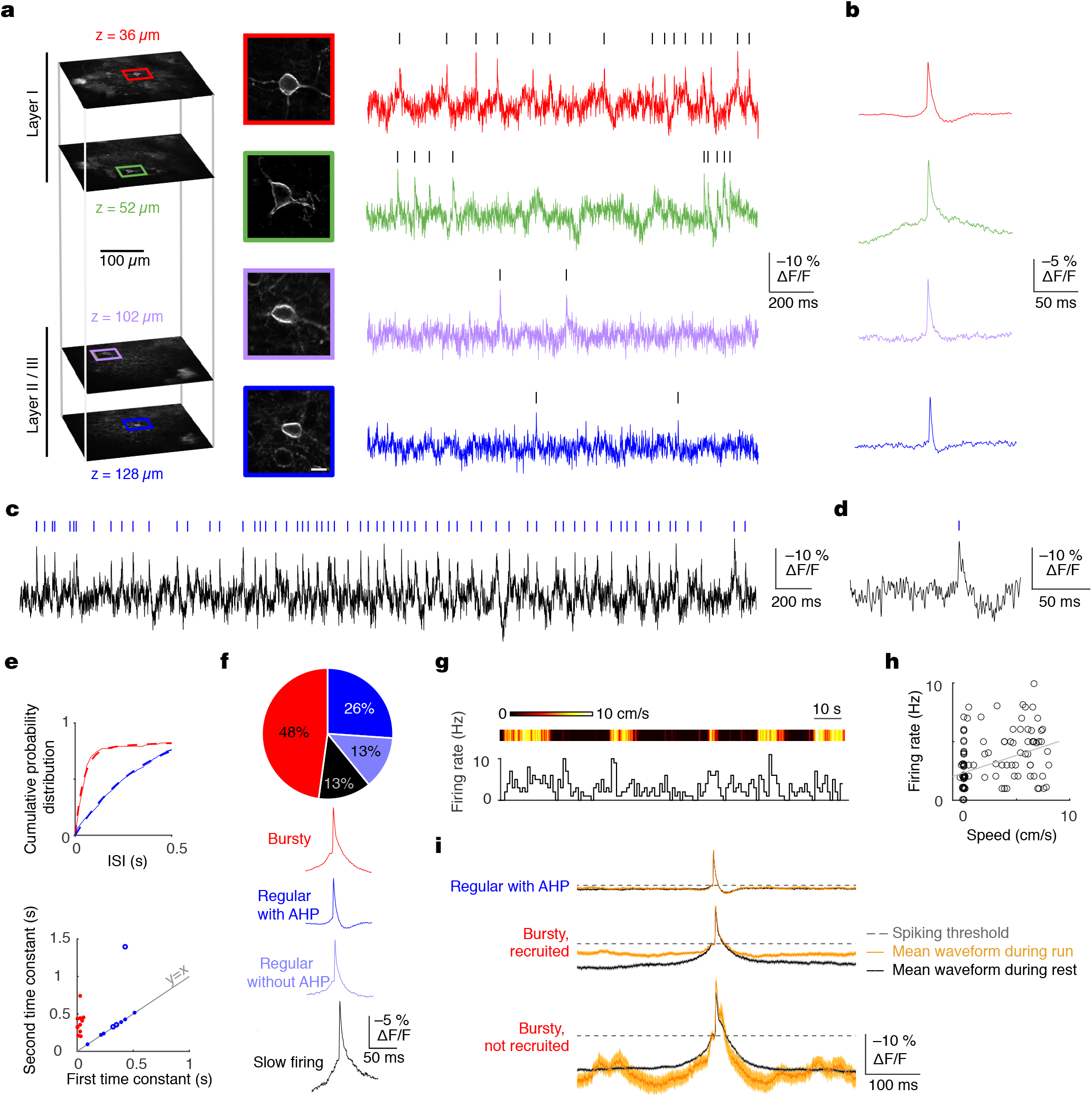
ASAP3-Kv reveals that modulation of bursting behavior by motor activity is cell type-dependent. **a**, Imaging ASAP3-Kv signals from neurons in different layers of one cortical column. **b**, Average AP shape reveals a variety of spike-related subthreshold behavior. **c**, Example of a recording from a regular-firing neuron with detected events (blue ticks). **d**, Zoomed-in example of a single spike with afterhyperpolarizing potential (AHP). **e**, Top, two representative cumulative inter-spike interval distributions with their tri-exponential fits (dashed lines). Bottom, scatter plot of the fitted time constant of the first and second components allows to segregate bursting neurons (red) from regular ring neurons (blue). **f**, Pie chart representing the classification of the 23 neurons analyzed in 4 groups with color coded average waveform for bursty neurons (11 cells, 2934 spikes, red), regular ring neurons with AHP (6 cells, 2405 spikes, blue), regular ring without AHP (3 cells, 803 spikes, light blue) and slow ring cells (3 cells, 77 spikes, black). **g**, Stairstep plot represents firing rate evolution together with animal running speed (heatmap). **h**, Relationship between firing rate and running speed (Pearson correlation coefficient r = 0.39, p < 0.0001). Gray line indicates the least-squares fit. **i**, Mean ± SEM. of the spike waveforms for cells during run periods (orange) and rest periods (black) for regular firing cells with AHP (4 cells, 827 and 1257 spikes during run and rest, respectively), bursty cells with ring rate increased by speed (3 cells, 601 and 411 spikes) or not increased (3 cells, 33 and 466 spikes). Dashed lines represent spike threshold chosen as a fixed membrane potential reference.

Next we examined differences in membrane potential dynamics surrounding APs during running vs. rest. We recorded five interneurons and six bursty pyramidal neurons during run episodes (**Fig. 6g-i**). Four of the interneurons and three of the pyramidal neurons were modulated positively by locomotion (for interneurons, Pearson correlation 0.355 ± 0.07, firing rate increase 3.1 Hz or 336 ± 341%, mean ± SD; for pyramidal neurons, Pearson correlation 0.306 ± 0.11, firing rate increase 1.63 Hz or 101 ± 58%). Interestingly, we found that the membrane potential of the interneurons in a 400-ms interval surrounding spikes was similar for spikes occurring during rest or locomotion, suggesting that spikes are emitted from the same depolarized subthreshold plateau potential in both conditions. In marked contrast, bursty pyramidal cells recruited by locomotion displayed a significantly more depolarized potential surrounding spikes during running than during rest (**Fig. 6i**, *∆F/F* = 3.6%, bootstrap p = 0.024). Interestingly, bursty pyramidal cells that were not recruited by locomotion showed the opposite effect of significantly more hyperpolarized potential surrounding spikes during running than during rest (**Fig. 6i**, *∆F/F* = 1.58%, bootstrap p < 0.001). Altogether, these results reveal that locomotor activity influences spiking rates in the visual cortex by regulating subthreshold dynamics surrounding spikes in a cell-type specific manner.

## DISCUSSION

In this study, we engineered an improved fluorescent GEVI, ASAP3, using a rapid library expression method and an automated electroporation-based screen, both developed *de novo* for this effort. ASAP3 features the largest brightness change across physiological voltages of any fluorescent indicator characterized so far (–50%), sub-millisecond activation kinetics to enable accurate AP timing, and extended deactivation kinetics to improve AP detection. Combining ASAP3 with RAMP microscopy and introducing a new method for ultrafast local volume excitation (ULOVE), we recorded transmembrane voltage of individual cortical and hippocampal neurons in awake head-fixed mice at sampling rates > 2 kHz, revealing oscillations and spikes in continuous recordings for many minutes. In this work, we demonstrate for the first time in the awake mammalian brain multiple abilities unique to the combination of genetically encoded voltage reporting and multi-photon microscopy: (1) single-cell single-trial voltage recordings in all cortical layers and in the hippocampus, (2) simultaneous voltage recordings from multiple dendrites, (3) optical recording of theta oscillations and spiking behavior in individual neurons, and (4) recording of voltage in the same neuron over multiple days.

The characteristics of ASAP3 and ULOVE help to overcome a long-recognized challenge for spike detection using GEVIs and point-scanning microscopy^6^. At the expression levels achievable with purely genetically encoded probes, which have well characterized maximal brightness relative to autofluorescence, a fluorescent transient of 1-ms duration would be difficult to separate from photon shot noise with high reliability^6^. Due to its 0.94-ms activation time constant at physiological temperature and large overall responsivity, ASAP3 responds to single APs with amplitudes beyond the previously presumed range^6^. ASAP3 also features a weighted deactivation time constant of 3.4 ms that prolongs the response while allowing discrimination of spikes in trains with frequencies up to 100 Hz, which covers the frequency range of most principal neurons^31^. In addition, ULOVE, by exciting reporter molecules over an extended 3D area of membrane, dramatically increases the available signal level and thus further enhances discriminability. As the ASAP3 response covers successive frames of fast ULOVE scanning and decays in an exponential manner, it can be recognized as distinct from noise by template fitting.

Our study also has implications for how GEVIs can be further improved. Historically, GEVI improvement has been a slow and laborious process due to lack of high-throughput screening methods and an incomplete understanding of GEVI sensing mechanisms. Our electroporation method induces voltage changes rapidly and robustly, and our approach of directly transfecting PCR reactions greatly accelerates library screening. PCR transfection has been independently developed^33–35^, but to our knowledge has not been used for improving genetically encoded indicators. As PCR transfection allows thousands of variants to be built and expressed in mammalian cells in a single day, this technique could be useful for improving other indicators that cannot be tested in bacteria. Our results also highlight the importance of structure-guided multi-site saturation mutagenesis, as both S150G and H151D mutations were required for the large improvement in ASAP3. These particular amino acid changes required three mutations at the DNA level, which, if random mutagenesis of the 1284-bp gene were performed to introduce exactly three mutations per clone, would arise once per 3.5 × 10^8^ (1284 choose 3) clones. Complete sampling of all 20 possible amino acids at two sites by random mutagenesis would be even less practical as it would require libraries with six mutations per gene, a complexity level of 6.2 × 10^15^ (1284 choose 6). Thus hypothesis-based saturation mutagenesis of multiple sites was crucial, adding to other examples of functional improvements via targeted multi-site mutagenesis^36–38^.

Interestingly, the maximal fluorescence change of ASAP3 across all voltages (4.5-fold) approaches that of GCaMP3 across all calcium concentrations^39^. Thus ASAP GEVIs, like GCaMP GECIs, are able to convert protein domain movements into large changes in the brightness of their circularly permuted GFP fluorophores. In addition, as with GCaMPs, only a small fraction of the fluorophore response range is utilized in response to a single spike, resulting in similar response amplitudes of ~20% for ASAP3 and current GCaMP variants^40^. If ASAP3 input sensitivity can be further narrowed to the physiological voltage range, larger responses to both subthreshold and spiking activity *in vivo* would be possible. Thus the identification of additional mutations that further tune ASAP3 input sensitivity to physiological voltages remains a priority.

In conclusion, the combination of a fast and responsive GEVI and kHz-rate ULOVE two-photon imaging enables investigation of neuronal information processing with a degree of specificity, resolution, and throughput that was previously not possible. ASAP3 yields sub-millisecond optical resolution of voltage spikes, an ability that calcium imaging lacks due to the inherently slower kinetics of calcium transients. ASAP3 also reports subthreshold membrane potential fluctuations, such as theta oscillations in hippocampal neurons or up and down states in layer-5 cortical neurons, events that are transparent to calcium indicators^41^. Recordings from brightly labeled isolated dendrites also opens the way for the optical study of synaptic integration in dendritic compartments inaccessible to electrodes, arguably a large missing component in our understanding of neuronal function. Taken together, our results demonstrate that theoretical advantages of GEVIs over electrodes in measuring transmembrane potential — genetic targeting, non-invasive access to deeply located neurons, parallel recording of multiple neurons, detection of subcellular voltage changes, and long-term recording *in vivo* — are now entering the realm of practice. We expect that ASAP3 and ULOVE two-photon microscopy will be broadly useful for high-speed recording of electrical events in genetically defined neurons in the brain.

## Methods

### Plasmid construction

Plasmids were made by standard techniques of molecular biology with all cloned fragments confirmed by sequencing. ASAP2s, subcloned into pcDNA3.1 with cytomegalovirus (CMV) promoter and bovine growth hormone terminator (BGHpA), served as the starting point for PCR-based library generation. For *in vitro* characterization in HEK293A cells, all ASAP variants and Arclight Q239 were subcloned in a pcDNA3.1/Puro-CAG vector^44^ between NheI and HindIII sites. For *in vitro* characterization in cultured neurons, and acute striatal slice experiments, ASAP variants were expressed from adeno-associated virus (AAV) packaging plasmids under the control of human synapsin promoter (hSYN). For kinetics characterization in CHO-K1 cells and soma-targeting in cortical slices, ASAPs were expressed from Cre-dependent plasmids (AAV-FLEX) downstream of EF1α promoter. To evaluate sensor brightness in neurons, ASAP variants were coexpressed with FusionRed-CAAX from the same AAV-hSYN plasmids through the encephalomyocarditis virus internal ribosome entry site (IRES). Farnesylation motif (CAAX) targeted FusionRed to the membrane to normalize for cell-to-cell differences in protein expression levels. For experiments in organotypic hippocampal slices, ASAP constructs were delivered to cells in pcDNA3.1/Puro-CAG vector plasmids. For acute striatal slice and mouse experiments, a soma-targeted version of the sensor, ASAP3-Kv, was created by fusing to the C-terminus of ASAP3 a linker of sequence GSSGSSGSS followed by the 65-aa proximal restriction and clustering signal from the C-terminal cytoplasmic segment of the Kv2.1 potassium channel^23^, which has been used to restrict opsins^45,46^ and ASAP2s^47^ to neuronal soma and proximal dendrites. ASAP3-Kv was then subcloned into pAAV-CAG-FLEX-WPRE (mouse) pAAV-hSYN-WPRE (slice), and the resulting plasmids were used to create AAV particles at the Neuroscience Gene Vector and Virus Core of Stanford University.

### Cell lines

All cell lines were maintained in a humidified incubator at 37°C with 5% CO_2_. HEK293A cells were cultured in high-glucose Dulbecco’s Modified Eagle Medium (DMEM, Life Technologies) supplemented with 5% fetal bovine serum (FBS, Life Technologies) and 2 mM glutamine (Sigma-Aldrich). The previously described HEK293-Kir2.1 cell line^12^ was further sub-cloned to achieve more consistent resting membrane potential of approximately –75 mV, and were maintained in high-glucose DMEM, 5% FBS, 2 mM glutamine, and 400 μg/mL geneticin (Life Technologies). For patch-clamp recordings HEK293A cells were transfected in 24-well plates with Lipofectamine 3000 (Life Technologies) per manufacturer’s recommended instructions (400 ng DNA, 1 μL P3000 reagent, 1 μL Lipofectamine). Between 4 and 24 hours after transfection, cells were re-plated at 15–25% confluency on 12-mm No. 0 uncoated glass coverslips (Carolina Biological), and were used the following day (1–2 days post-transfection). For electrical screening HEK293-Kir2.1 cells were plated in 384-well plates (Grace Bio-Labs) on conductive glass slides coated with poly-D-lysine hydrobromide (Sigma-Aldrich). Glass slides were conductive due to one-sided coating with indium tin oxide with surface resistance of 70–100 Ω (Delta Technologies). HEK293-Kir2.1 cells were transfected in 384-well plates with Lipofectamine 3000 (~100 ng DNA, 0.4 μL p3000 reagent, 0.4 μL Lipofectamine) followed by a media change 4–12 hours later, and imaged 2 days post-transfection. For characterization of GEVI kinetics at physiological temperatures, CHO-K1 cells (ECACC) were cultured in DMEM:F12 with 10% FBS, and were transfected with ASAP constructs by polyethylenimine (PEI).^45^

### Library construction and expression

Libraries of linear ASAP variants were constructed in deterministic manner. To enable transient expression of ASAP variants in mammalian cells, libraries of linear ASAP constructs were designed to include the cytomegalovirus (CMV) promoter and the bovine growth hormone polyadenylation signal sequence (BGHpA). Libraries of variants were generated by high fidelity PCR amplifications with PrimeSTAR HS DNA Polymerase (Takara) or Phusion Flash II DNA Polymerase (Thermo Fisher Scientific). Synthetic DNA oligonucleotides were purchased from Integrated DNA Technologies or Eurofins Genomics. For libraries concurrently targeting 2 amino acids, each of the possible 400 (20 × 20) combinations of mutations were carried by a single primer. Libraries of primers were ordered in 384-well plate format, and were used to generate final constructs by overlap extension PCR from two templates. The first template contained CMV promoter and an unchanged part of ASAP (N-terminal region and S1 to S3), while the second template included the remainder of ASAP carrying combinatorial mutations and BGHpA. For any single library, the first PCR template was constant across all constructs, and was purified by preparative agarose gel electrophoresis and Zymoclean gel recovery kit (Zymo Research). The second template, generated by mutagenizing primers, was diluted 100-fold and used in subsequent overlap extension PCR without any cleanup. Approximately 100 ng/μL of final PCR products were typically generated during the final PCR reaction (30 PCR cycles), and were directly used for transient transfection in 384 well-plates without any cleanup. During all steps of PCR assembly select wells from 384-well plates were monitored by gel electrophoresis. For the final product size of only 2216 bp, the estimated percentage of PCR products with an error is 6.32% when using high fidelity Phusion Flash PCR Master Mix (error rate of 9.5 × 10^-7^). Small libraries in which saturating mutagenesis was performed at a single site were generated in analogous way. PCR-generated linear constructs were stable if kept at –80 °C, and were typically used for multiple transfection rounds. Transfection efficiency with PCR products kept at –20 °C rapidly decreased even within several days.

### Electrical screening in HEK293-Kir2.1 cells

For functional screening by electroporation we imaged cells on an IX81 inverted microscope fitted with a 20× 0.75-numerical apertures (NA) objective (Olympus). A 120-W Mercury vapor short arc lamp (X-Cite 120PC, Exfo) served as the excitation light source. Filter cube set consisted a 480/40-nm excitation filter and a 503-nm long pass emission filter. Cells were plated in 384-well plates on conductive glass slides, and imaged directly on the microscope stage in Hank’s Balanced Salt solution (HBSS) buffered with 10 mM HEPES (Life Technologies). Unless mentioned otherwise, the conductive glass slide was connected to the ground terminal of an S48 Grass stimulator (Astro-Med). The electrical circuit was completed by a platinum electrode (0.25 mm in diameter, Sigma-Aldrich) brought into each well and placed ~ 400 μm above the monolayer of cells. A custom-built holder, secured in place of the microscope condenser, supported the assembly for platinum electrode movement in z-dimension and its connectivity to the stimulator. ASAP libraries were screened on the platform in a semi-automated mode with the operator locating the best field of view and focusing on the cells, although the system is capable of a fully automated mode as well. A single field of view was imaged for a total of 5 s, with a square 10-μs 150-V square pulse applied at the ~3 s mark. Fluorescence was recorded at 100 Hz (10-ms exposure per frame) by an ORCA Flash4.0 V2 C11440-22CA CMOS camera (Hamamatsu) with pixel binning set to 4 × 4. Image acquisition, stage advancement from well-to-well, electrode positioning and pulse application were controlled by MATLAB 9.0 (Mathworks). We utilized an automated image analysis routine to determine the average of a maximum change in fluorescence in a single field of view. The program bins images such that a single pixel corresponds to approximately one cell. The region with the lowest fluorescence serves as the background correction for the signal. The change in fluorescence following the pulse is determined for all such cells. Five cells with the largest change in signal are selected and an average of their responses is used in analysis. Full combinatorial libraries were screened at least three times, with the top 40 variants further characterized on a secondary screen at 4–6 wells per variant. If there were no better variants than the parent (screening round 2), no secondary round was performed. To determine which variants to patch, data across all screening runs was evaluated. This way each construct was measured at least three times (from primary screens) or more (if included in the secondary screen).

### Primary cell culture

Hippocampal neurons were isolated from embryonic day 18 Sprague Dawley rat embryos by dissociation in RPMI medium containing 5 units/mL papain (Worthington Biochemical) and 0.005% DNase I at 37 °C and 5% CO_2_ in air. Dissociated neurons were plated at a density of 3.5 × 10^4^ cells/cm^2^ in 24-well plates on washed 12-mm No. 0 glass coverslips pre-coated overnight with > 300-kDa poly-D-lysine hydrobromide (Sigma-Aldrich). Cells were plated for several hours in Neurobasal media with 10% FBS, 2 mM GlutaMAX, and B27 supplement (Life Technologies), then media were replaced with Neurobasal with 1% FBS, 2 mM GlutaMAX, and B27. Half of the media was replaced every 3–4 days with fresh media without FBS. 5-Fluoro-2′-deoxyuridine (Sigma-Aldrich) was typically added at a final concentration of 16 μM at 7–9 DIV to limit glia growth. Neurons were transfected at 9–11 DIV using a modified Lipofectamine 2000 (Life Technologies) transfection procedure in which media in one well of a 24-well plate was replaced for 60–90 min with 200 μL of DNA-lipid complexes (100 ng indicator DNA, 400 ng empty pNCS vector, 1 μL Lipofectamine 2000, 200 μL Neurobasal with 2 mM GlutaMAX). Procedures were carried out in compliance with the rules of the Stanford University Administrative Panel on Laboratory Animal Care.

### Whole cell patch clamping and imaging of HEK293A cells

Patch-clamp experiments were mainly done as previously described^11^. Electrical signals were recorded in voltage-clamp mode with Multiclamp 700B amplifier and pClamp software (Molecular Devices). 3.5- to 5-MΩ patch electrodes were pulled from borosilicate capillaries with filament and 1/1.5 mm ID/OD (King Precision Glass). HEK293A cells, sparsely plated on uncoated glass coverslips, were kept in a perfusion chamber on the stage of Axiovert 100M inverted microscope (Zeiss), and were continuously perfused with extracellular solution (110 mM NaCl, 26 mM sucrose, 23 mM glucose, 5 mM HEPES-Na, 5 mM KCl, 2.5 mM CaCl_2_, 1.3 mM MgSO_4_, pH adjusted to 7.4) at room temperature (23 °C). Patch pipettes were filled with 115 mM potassium gluconate, 10 mM HEPES-Na, 10 mM EGTA, 10 mM glucose, 8 mM KCl, 5 mM MgCl_2_ and 1 mM CaCl_2_, pH adjusted to 7.4. Fluorophores were illuminated at ~ 4.3 mW/mm^2^ power density at the sample plane with an UHP-Mic-LED-460 blue LED (Pryzmatix) passed through a 484/15-nm excitation filter and focused on the sample through a 40× 1.3-NA oil-immersion objective (Zeiss). Emitted fluorescence passed through a 525/50-nm emission filter and was captured by an iXon 860 electron-multiplied charge-coupled device camera (Oxford Instruments) cooled to –80 °C. For all experiments Frame Transfer mode was enabled, and EM gain was set to 10. To characterize steady-state fluorescence responses step voltage commands (500-ms duration) were applied from a holding potential of –70 mV to voltages ranging from –120 to 50 mV for standard F–V curves, and to depolarizing (–70 to +180 mV) or hyperpolarizing (0 to –180 mV) voltage steps for F–V curves over extended voltages. When voltage-clamp could not be imposed on some cells at the highest potentials, that step was excluded from data analysis. A 10-ms depolarizing pulse (from –70 to –16 mV) was included before each step to simplify image processing and allow coarse assessment of sensor kinetics. Images were captured at 200 Hz, and fluorescence was analyzed from the perimeter of the cell. To characterize kinetics of ASAP indicators, images were acquired at 2.5 kHz from an area cropped down to 64 × 64 pixels and further binned to comprise only 8×8 pixels. Fluorescence was quantified by summing signal from all 64 pixels. Command voltage steps were applied for 1 s, and an average of two identical steps was contributed by each cell for analysis. Double exponential fits were applied to a 500-ms interval after the voltage step using IgorPro sofware (WaveMetrics). For all experiments fluorescence traces were corrected for photobleaching by dividing the recorded signal by an exponential fit to the data. Data were fit using a double exponential or a single exponential (once the initial rapid photobleaching was excluded). Only HEK293A cells with access resistance (*Ra*) < 12 MΩ and membrane resistance (*Rm*) to *Ra* ratio > 20 were included in the analysis. Command waveforms imposed in voltage clamp accounted for the liquid junction potential.

### Whole cell patch clamping and imaging of CHO-K1 cells

Transient expression of ASAP2 and ASAP3 in CHO-K1 cells was achieved by PEI transfection^48^. Constructs were driven by insertion of the GEVI cDNAs into an AAV expression plasmid downstream of an eF1α promoter, and flanked by pairs of loxP and lox2272 sites for cre-dependent expression. Cells were transfected with a 200-μL mixture of PEI and DNA (pAAV-EF1α-ASAP and pBS185-CMV-CREplasmids^49^). PEI (25 kD at 1 mg/mL), was mixed at 3:1 ratio (w/w) with 1.5 μg of GEVI and 0.5 μg of CRE plasmids, and added to cells two days after plating on 15-mm sterile glass coverslips in a well of a 6 well plate. Whole-cell patch mode recordings were performed at 22 or 34 °C after 3–4 days of expression. External solution was 140 mM NaCl, 2.8 mM KCl, 1 mM CaCl_2_, 10 mM HEPES pH 7.3. Internal solution was 10 mM EGTA, 0.2 mM CaCl_2_, 110 mM CsCl, 20 mM TEA, 10 mM HEPES, 6 mM NaCL, 4 mM ATP, 0.4 mM GTP. Voltage-clamp recordings from CHO-K1 cells were performed with a Multiclamp 700B amplifier, filtered at 10 kHz, and then digitized at 50 kHz. Series resistance compensation of CHO cells resulted in a final series resistance below 5 MΩ. Fluorescence imaging was performed on a BX51 upright microscope (Olympus) equipped with platforms and manipulators (Scientifica) and a Xzyla camera (Andor). ASAP excitation was obtained by restricted illumination of the isolated recorded cell using a SOLA light engine (Lumencor) with the following filters: excitation ET470/40X, dichroic T495LPXR, emission ET525/50M (Chroma). The fluorescence signal was acquired by a PMTSS photomultiplier tube (Thor Labs) low-pass filtered at 3 kHz and digitized at 50 kHz.

### Whole cell patch clamping and imaging of cultured hippocampal neurons

Cultured rat hippocampal neurons were used for patch-clamping at 11–14 DIV, 2–4 days post-transfection. Patch-clamp recordings were performed as above with HEK293A cells. Bath and intracellular solutions were the same, except CNQX disodium salt hydrate (Sigma-Aldrich) was added to the extracellular solution in some cases to isolate current-evoked APs. Recordings in neurons were performed in current-clamp mode without holding current injection. APs were elicited by injecting 1-ms pulses of 700–1100 pA of current. pClamp (Molecular Devices) software was used to record electrode voltages, which were corrected for the liquid junction potential post hoc. Images were captured at 1 kHz with 4 × 4 binning, and fluorescence was measured from all pixels of the cell body. Only neurons with resting membrane potential less than –50 mV, *Ra* less than 17 MΩ, and *Rm* at least 10-fold greater than *Ra* were analyzed. For those cells, only APs with a peak amplitude of at least 0 mV and width less than 5 ms at –20 mV were used for ASAP characterization. Three to 34 APs were captured per neuron, and their average (APs per cell) was used for statistics. Fluorescence traces were corrected for photobleaching and normalized to baseline by custom routines in MATLAB, then exported to Excel (Microsoft) for graphing. For **Supplementary Movie 1** and **Supplementary Fig. 8a**, ASAP3-expressing hippocampal neurons were imaged while in voltage-clamp mode with an ORCA Flash4.0 V2 C11440-22CA CMOS camera (Hamamatsu) at 200 Hz.

### Evaluation of brightness and soma-targeting in hippocampal neurons

GEVIs were transiently expressed in cultured hippocampal neurons and imaged 2 days plater at 12 DIV in extracellular solution. FusionRed with a C-terminal farnesylation motif for membrane targeting was co-expressed from the same plasmids via an internal ribosome entry site. Cells were imaged on an Axiovert 200M inverted microscope with a 20× 0.75-NA (for brightness) or 40× 1.2-NA (soma-targeting) objective (Zeiss) and an X-Cite 120 metal-halide lamp (Lumen Dynamics) as the excitation light source. The following excitation (ex) and emission (em) filters were used: ASAP, ex 490/10-nm, em 525/40-nm; FusionRed, ex 555/25-nm, em 620/60-nm. Images were captured unbinned on an Orca-ER CCD camera (Hamamatsu) with Micro-Manager software^50^. Image processing was done using a custom algorithm written in MATLAB that segmented neurons into soma and neurite regions and quantified fluorescence from the perimeter for both channels. ASAP fluorescence intensity was then divided by FusionRed intensity to obtain a measure of relative brightness per molecule or to determine depletion of ASAP3-Kv signal in neurites relative to the soma.

### Confocal imaging of cortical slices

Stereotaxic injections of ASAP3 and ASAP3-Kv AAVs were performed on C57BL/6 mice under ketamine/xylazine anesthesia. AAV2/1-EF1α-ASAP3 or AAV2/1-EF1α-ASAP3-Kv (obtained from IBENS core, both at 3.5 × 10^12^ genome copies (GC)/mL) were diluted 1:5, and co-injected with AAV2/1-hSYN-Cre (Penn vector core, 1.9 × 10^13^ GC/mL, 1:500 dilution). 500 nL of virus mixtures were injected at 200 nL/min into the same animal, with ASAP3 into the left hemisphere (coordinates AP 2.5 mm, ML 2.5 mm, DV 0.33 mm) and soma-targeted ASAP3-Kv into the right (AP 2.5 mm, ML –2.5 mm, DV 0.33 mm). After 20 days mice were transcardially perfused with 4% Antigenfix (Diapath), and brain post-fixed overnight in Antigenfix, followed by PBS for five hours. 50-μm thick slices were cut on a VT100S vibratome (Leica) and mounted on slides between coverslips in Vectashield (Vector Labs). Images were captured using LSM 510 (Zeiss) and displayed as maximal projection of confocal z-stacks.

### Patch-clamp electrophysiology and two-photon imaging in acute striatal slices

ASAP3-Kv expression was achieved via adeno-associated virus. Stereotaxic injections were performed on P40-P60 C57BL/6 mice under ketamine anesthesia. 900 nL of 3.95 × 10^13^ GC/mL AAV9-syn-ASAP3b-Kv virus was injected unilaterally (right hemisphere) into the dorsolateral striatum (at AP 1.2 mm, ML –2 mm, and DV –3.2 mm from bregma). Injection was performed using a micropipette (VWR) pulled with a long narrow tip (size ~10–20 μm) using a micropipette puller (Sutter Instrument). The glass micropipette was inserted into the brain and virus was injected at an infusion rate of 200 nL/min. The pipette was gently withdrawn 5 min after the end of infusion and the scalp was sutured. Animals were used at 4–12 weeks after AAV injections. Coronal brain slices (300 μm) containing the dorsal striatum were obtained using standard techniques^51^. Briefly, animals were anesthetized with isoflurane and decapitated. The brain was exposed, chilled with ice-cold artificial CSF (ACSF) containing 125 mM NaCl, 2.5 mM KCl, 2 mM CaCl_2_, 1.25 mM NaH2PO4, 1 mM MgCl_2_, 25 mM NaHCO3, and 15 mM D-glucose (300–305 mOsm). Brain slices were prepared with a vibrating microtome (Leica VT1200 S, Germany) and left to recover in ACSF at 34°C for 30 min followed by room temperature (20–22 °C) incubation for at least additional 30 min before transfer to a recording chamber. The slices were recorded within 5 hours after recovery. All solutions were saturated with 95% O_2_ and 5% CO_2_. Striatum MSNs were visualized under infrared illumination using an Olympus BX-51 microscope equipped with DIC optics, a water-immersion objective (60× NA 1.0). ASAP3-Kv expressing neurons were identified under epifluorescence illumination (Lambda XL, Sutter Instrument). Whole-cell current-clamp recording was performed with borosilicate glass microelectrodes (3-5 MΩ) filled with a K+-based internal solution (135 mM KCH_3_SO_3_, 8.1 mM KCl, 10 mM HEPES, 8 mM Na_2_-phosphocreatine, 0.3 mM Na_2_GTP, 4 mM MgATP, 0.1 mM CaCl_2_, 1 mM EGTA, pH 7.2–7.3, 285-290 mOsm). Access resistance was < 25MΩ and compensated by applying bridge balance. To induce MSN firing, 2-ms pulses of 2-nA current were injected to induce spiking at different frequencies (10, 50, 100, or 200 Hz). Recordings were obtained with a Multiclamp 700B amplifier (Molecular Devices) using the WinWCP software (University of Strathclyde, UK). Signals were filtered with a Bessel filter at 2 kHz to eliminate high frequency noise, digitized at 10 kHz (NI PCIe-6259, National Instruments). Two-photon imaging was performed with a custom built two-photon laser-scanning microscope as described previously^52^ equipped with a mode-locked tunable (690–1040 nm) Mai Tai eHP DS Ti:sapphire laser (Spectra-Physics, USA) tuned to 920 nm. ASAP3-Kv signals were acquired by a 1-kHz line scan across the membrane region of a cell. Signals recorded along each line were integrated to produce fluorescence trace over time.

### Patch-clamp electrophysiology in cultured hippocampal slices

Preparation and culture of organotypic hippocampal slices was done as previously described^11^. Organotypic hippocampal slices were carefully laid flat and immobile with a nylon mesh in an electrophysiology chamber. During experiments, the slices were continuously perfused with an oxygenated (mixture of 95% O_2_ and 5% CO_2_) solution of articifial cerebrospinal fluid (ACSF) containing 124 mM NaCl, 25 mM NaHCO_3_, 2.5 mM KCl, 1.2 mM MgCl_2_, 2.5 mM CaCl_2_, 10 mM glucose (pH = 7.4, 300 mOsm). The ACSF was warmed to a temperature of 32 ± 2°C as measured in the recording chamber or was kept at room temperature which resulted in ACSF at 20 ± 2°C in the recording chamber. A Dodt scanning gradient contrast was used to guide the pipette relative to the slice for patch-clamp recordings. ASAP-expressing neurons were identified under two-photon illumination. The pipette (resistance in the bath of 3–5 MΩ) was carefully approached to targeted neurons. The intracellular solution contained 120 mM K-gluconate, 20 mM KCl, 10 mM HEPES, 2 mM MgCl_2_, 2 mM Mg_2_ATP, 0.3 mM NaGTP, 7 mM phosphocreatine, 0.6 mM EGTA (pH = 7.2, 295 mOsm). Slight negative pressure was applied to break the giga-seal to the whole-cell configuration. All experiments were performed in the current-clamp mode. The electrophysiological signal was amplified with a Multiclamp 700B (Molecular Devices) and low-pass filtered at 2 kHz. The signal was digitized at 10 kHz with a Digidata 1440A (Molecular Devices) and recorded with Clampex 11.0 software (Molecular Devices). Action potentials were evoked by 2- to 4-ms square current pulses (1.0–1.5 nA). Trials were acquired at 10-s intervals.

### Patch-clamp electrophysiology in acute cerebellar slices

Animals were anesthetized with isoflurane (4% in medical oxygen) and immediately decapitated. The cerebellum was rapidly removed and placed in an ice-cold BBS solution (125.7 mM NaCl, 3.3 mM KCl, 1.25 mM NaH_2_PO_4_, 24.8 mM NaHCO_3_, 25 mM glucose, 1.3 mM CaCl_2_, 1.17 mM MgCl_2_, 50 nM minocycline). The cerebellum was glued (Cyanolit) in the slicing chamber on a parasagittal section and submerged in ice-cold cutting solution (130 mM K-gluconate, 14.6 mM KCl, 2 mM EGTA, 20 mM HEPES, 25 mM glucose, 50 μM D-APV, 50 nM minocycline) during slicing. Slices of 290-μm thickness were cut using a stainless steel blade (z-axis deflection < 0.5 μm) with an oscillating blade microtome (Campden Instruments) and kept in warm (33°C) oxygenated recovery solution (225 mM D-mannitol, 2.3 mM KCl, 1.25 mM NaH_2_PO_4_, 25 mM NaHCO_3_, 25 mM glucose, 0.51 mM CaCl_2_, 7.7 mM MgCl_2_, 50 μM D-APV, 50 nM minocycline) for several minutes before being transferred to a chamber recirculated with warm (33°C) oxygenated BBS solution. At least 30 min after being cut, slices were transferred to a recording chamber mounted on the AOD-based multiphoton microscope. Slices were perfused with 3.5 mL/min gassed BBS solution at 33 °C. Borosilicate glass patch electrodes (resistance 4–6 MΩ) were filled with a high-chloride intracellular solution (135 mM KMeSO_4_, 6 mM NaCl, 2 mM MgCl_2_, 10 mM HEPES, 4 mM ATP-Mg, 0.4 mM GTP-Na_2_, adjusted at pH 7.35 with KOH (290–295 mOsm). Whole-cell patch-clamp recordings were performed using a Multiclamp700B amplifier, Digidata digitizer, and PClamp10 software (Molecular Devices). Molecular layer interneurons of the cerebellar cortex were held around –70 mV in the whole-cell current-clamp configuration and large currents (1 ms, 300–800 pA) were injected to trigger a spike. Electrophysiological signals were digitized at a sampling rate of 20 kHz and passed through a 5-kHz Bessel filter.

### Random-access two-photon voltage imaging in hippocampal slices

A custom-built random-access two-photon microscope (RAMP) was used to perform voltage imaging^53^. A Chameleon Ultra II titanium:sapphire laser (Coherent) with a repetition rate of 80 MHz (140 fs, average power > 4 W) was tuned at 900 nm. The laser beam passed through an acousto-optic modulator, for spatial and temporal chromatic compensation, and on a pair of AODs (A-A Opto-Electronics) to rapidly direct the laser beam in x and y dimensions in the field of view. The laser beam was relayed through a 4f system from the middle plane of the x-y AOD scanner to the back-aperture of a 25× 0.95-NA objective (Leica). Photons were collected in either the transfluorescence mode (through the condenser), or in both the trans-fluorescence and epifluorescence modes simultaneously. Both detection pathways were the same. The signal was passed through a 720-nm short-pass filter and then split in two channels using a 580-nm dichroic mirror (Semrock), resulting in a green (ASAPs) and a red channel (AlexaFluor-594). Photons in the green channel were band-pass filtered at 500–560 nm and photons in the red channel were band-pass filtered at 595–665 nm (Semrock). Photons were detected using H7422P-40 AsGaP external photomultiplier tubes (Hamamatsu) in the photon-counting mode. Photon counts were acquired on a personal computer and the microscope was controlled with a software written in LabVIEW.

### ASAP expression in mice

Male adult wild-type C57BL/6 mice (n = 9, 22–28 g body weight) were used in the experiments. All protocols adhered to the guidelines of the French National Ethic Committee for Sciences and Health report on ‘‘Ethical Principles for Animal Experimentation’’ in agreement with the European Community Directive 86/609/EEC under agreement #02 235.02. All mice were housed in standard conditions (12-hour light/dark cycles light on at 7 a.m., housed two or three per cage, with water and food ad libitum). AAV2/1.Syn.Cre was obtained from the Penn Vector Core at a concentration of 1.9 × 10^13^ GC/mL. AAV2/9.CAG.Flex.ASAP3-Kv was produced by the Stanford Neuroscience Gene Vector and Virus Core facility at a concentration of 2.05 × 10^13^ GC/mL. AAV2/1.Syn.Cre or AAV2/1.CamKII.Cre diluted 1:1000 and AAV2/9.CAG.Flex.ASAP3-Kv diluted 1:5 to 1:100 were combined into phosphate-buffered saline (PBS), 400 nL of which was injected at a flow of 50 nL/min into the cerebellum, the CA1 area of the hippocampus and sensory cortices (CA1 coordinates from bregma: AP –2.3 mm, ML –2 mm, DV –1.3 mm from dura, n = 2 mice; S1FL: AP –0.3 mm, ML –2.5 mm, DV –0.3, n = 1 mouse; V1: AP –3 mm, ML –2.5 mm, DV –0.3 mm). A preoperative analgesic was used (buprenorphine, 0.1 mg/kg). Hippocampal surgery was performed 7 days later as previously described^26,54^, but without water restriction. Briefly, a glass-bottomed cannula was inserted on top of the dorsal hippocampus after aspiration of the cortex, and secured with Kwik-Sil at the tissue interface and Superbond at the skull level. For the primary sensory and visual cortices, a 3- or 5-mm coverslip was placed on top of the targeted cortical area immediately after viral injection and secured with VetBond and dental cement. For both surgeries, a custom designed aluminum headplate was fixed on the skull with layers of dental cement after the coverslip implantation. Mice were allowed to recover for at least 15 days before imaging sessions.

### Two-photon voltage imaging by ultrafast local volume scanning in cerebellar slices and awake mice

Multiple imaging sessions of 1–3 hours duration over 1–4 days were performed while mice behaved spontaneously on a running wheel in the dark. Behavioral habituation involved progressive handling by the experimenter with gradual increases in head fixation duration^26^. Mice were handled before recording sessions to limit restraint-associated stress and experiments were performed during the light cycle. Imaging was performed using a custom designed AOD-based random-access multi-photon system based on a previously described design^53^ (Karthala System) and a Ti:sapphire femtosecond laser (Spectra Physics) mode-locked at 920 nm with a repetition rate of 80 MHz. A 25× water-immersion objective (0.95-NA, 2.5-mm working distance, Leica) was used for excitation and epifluorescence light collection. The signal was passed through a 720-nm shortpass filter, split into two channels using a 580-nm dichroic mirror (Semrock, New York, USA), and passed to two H10769/40 cooled photomultiplier tubes (Hamamatsu) in photon counting mode, with the green channel used for ASAP3. For acute slices the trans-fluorescence signal was similarly split into two channels using a 580-nm dichroic mirror, collected by two H10769/40 cooled photomultiplier tubes in photon counting mode, and summed with the epifluorescence signal. An AOD-based local volume scanning method was used to achieve high sampling rate (3–5 kHz) and stable recordings *in vivo* (Mathieu et al., manuscript in preparation). In short, each voxel consisted in a parallelepiped of about 3 × 6 × 6 μm scanned within 100 μs. All fluorescence in this volume was integrated to yield one recording value. For local volume scanning non-stationary patterns of frequencies were digitally encoded and fed to a DDS device at a rate of 10 MHz. The frequency modulated output signal was amplified and drove the AOD scanners. Frequency patterns consisted in linear chirps and sinusoidal modulations, which could be varied in amplitude to tune the dimension of the local volume. An HEDS-5645#A06 optical quadrature encoder (Avago Technology) was used to keep track of instantaneous speed of the mouse on the running wheel with a spatial resolution of 0.484 mm.

### ASAP3 signal analysis

Due to the diversity of spike shapes (presence of after-depolarization or after-hyperpolarization) and firing properties (high burstiness or regular firing) in our sample, we had to implement a unique detection method consisting of three successive steps. First, we isolated the biggest events to generate a realistic waveform template. Second, we further detected isolated events based on correlation with this template. This second step generates a more precise waveform, yielding parameters to feed MLspike^24^. Third, we performed the final detection using MLspike for template peeling in bursts. For each cell the ASAP3 signal was obtained by pooling the fluorescence from 2 or 3 volumes of interest encompassing different regions of the cell membrane. Slow drift was removed from the obtained recording by subtracting a bi-directional low-pass filtered trace (< 20 Hz, 4th-order Butterworth filter) and rare periods with shivering related motion artefacts were discarded. We then computed a local z score of the trace, and events crossing a threshold of 4 were detected. During the second step, the average of the detected events was used as a template for a template-matching correlation method. The temporal extent of the template was defined by an iterative method where correlation histogram skewness was maximized. A local maximum version of the correlation histogram was computed with a window corresponding to the template duration. If its density distribution was bimodal its local minimum was used as a threshold, else the whole recording was discarded. This method detects all isolated spikes but not all spikes within bursts, due to template length. The decay of the resulting averaged spike waveform was then fitted by a bi-exponential function to extract the average amplitude and decay time constant of the optical spikes. Finally, we fed a model-based spike inference algorithm in MLspike with the extracted amplitude and decay tau, in order to isolate both isolated spikes and spikes within bursts. For each neuron, a final average spike waveform was then obtained and the amplitude (*∆F/F*) was measured as the fractional decrease of fluorescence at the peak. For this a local baseline value was taken as the average of the signal for the 3 ms preceding the spike occurrence in MLspike. To limit shot noise influence on single-point local maximum, single event peak value was obtained by averaging the signal over 2 ms following spike detection and correcting for the exponential decay of the spike according to the corrective factor

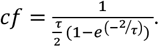

The discriminability index d’ was calculated using the published formula 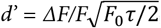, where *F*_0_ is the photon flux and *τ* is the spike duration taken as the decay time constant of the average waveform^22^. Both the Gaussian shape of the event amplitudes distribution, which was not truncated **(Fig. 4g)**, and the high discriminability index of the analyzed cells **(Fig. 4f)** indicated that the false negative rate was small. Mean firing rate was defined as the number of detected deflections divided by the duration of the analyzed period (typically 150 s). In a first evaluation, bursts were defined as a succession of deflections separated by a maximal inter-spike interval of 25 ms and burstiness as the fraction (expressed in percentage) of spikes within bursts. In order to classify cells more precisely, we fitted the interspike interval cumulative distribution with a tri-exponential fit. All neurons displayed at least two temporal components, but neurons were considered as bursty if all three time constants were significantly different, indicating the existence of a third component of short interspike intervals. When goodness of fit was smaller than 1 because of the small number of spikes, cells were assigned to a slow firing group.

We further classified neurons based on their membrane potential dynamics following the spike. A smoothed differential signal was computed by subtracting the averages over two 3-ms sliding windows separated by 8 ms. We then detected the points of this differential signal in a 20- to 50-ms time window following the spike which departed significantly from zero, as assessed from baseline noise. Applying this to our cell sample revealed that 6 cells of the regular firing group displayed a significant after-hyperpolarizing potential, while all other cells displayed a significant after-depolarizing potential.

To evaluate if cells were recruited during locomotion, we first obtained the instantaneous position of the mouse from the optical encoder in the running wheel^26^. A firing rate and a speed timecourse were calculated by averaging over 1-s bins. Cells with at least one bin above 4 cm/s were retained for this analysis. The significance of the linear correlation between the firing rate and speed vectors was used to establish recruitment by locomotion. Locomotion and rest periods were defined using previously published criteria^26^, and the average spike shape for those two behavioral periods was pooled over cells within one cell group. To test for the significance of the influence of locomotion recruitment on membrane potential dynamics within and between cell classes, we performed a bootstrap with 1000 surrogates taken randomly amongst all cells.

To quantify theta oscillations, a Morlet wavelet-based spectrogram was used to obtain the instantaneous power, frequency and phase of the hippocampal theta oscillations (5–10 Hz). Run periods were extracted with a previously validated algorithm^26^, consisting of epochs where the mouse moves forward for at least two centimeters at a speed above two cm/s. The power profile is the average of the spectrogram for a given behavioral state across frequencies. For speed/theta correlation analysis, the animal’s speed was binned every 0.5 cm/s. For optical spikes/theta oscillation phase-locking, we looked for the distribution of optical spike times on the oscillation phase at the peak frequency with 2π/16 radians bin size. We normalized the distribution by the number of deflections per cell. For display purposes only, two cycles are shown.

### Statistical analysis

For differences between ASAP2s and ASAP3 in responses to APs in cultured neurons, a power calculation was performed in the G*Power 3 program^55^ to determine the number of neurons required for detecting a 25% difference in response amplitudes at an alpha level of 0.05, based on measured variability in ASAP2s response amplitudes^11^. For differences between ASAP variants in brightness in cultured neurons, variability of brightness was measured in a pilot experiment and a power calculation was performed in G*Power 3 to determine the number of neurons required for detecting a 25% increase in brightness between any two indicators among five indicator variants at an alpha level of 0.05. Brightness was then measured in ≥ 35 neurons. Normality of the data were confirmed by F-test. One-way ANOVA followed by Tukey’s posthoc test was performed. For differences between ASAP variants in response amplitude and kinetics in organotypic slices, the number of samples required to detect a difference of 50% in response amplitude or brightness at an alpha level of 0.05, given variability in response amplitudes and half-decay times previously observed in the same experimental system^11^ was calculated in G*Power 3. Normality of the data were confirmed by F-test and dependent variables were tested using a two-sided t-test. For differences in ASAP3 responses between cell populations *in vivo*, dependent variables were tested using the Wilcoxon rank sum test. Sample numbers were dictated experimentally by the number of cells that could be imaged by one experimenter within a week, to enable cells to be approximately matched in mouse age.

## Code and data availability

All custom code and data from this study are available from the corresponding authors upon request.

## Acknowledgements

We thank Lin Ning (Lin laboratory), Luke Kaplan, and Bianxiao Cui (Stanford University) for assistance with neuronal cultures, Chaleen Salesse and Paul De Koninck (CERVO Brain Research Centre, Université Laval) for assistance with organotypic slice cultures, Patrick Houlihan and David Clapham for advice on CHO cell culture and transfection, Stephane Supplisson for assistance with electrophysiology on CHO cells, Annick Ayon for assistance with ASAP subcloning and viral vector constructions and Walther Akemann, Astou Tangara and Yvon Cabirou (ENS facilities, Paris) for assistance with *in vivo* imaging. We also thank Gui-Rong Li (University of Hong Kong) for the gift of the HEK293-Kir2.1 cell line and Karl Diesseroth (Stanford University) for the gift of pAAV-CAG-FLEX-WPRE plasmid. This work was supported by the Marc A. Cohen Graduate Fellowship Research Fund, the Stanford Bioengineering PhD Program and American Epilepsy Society predoctoral fellowship (M.C.); the China Scholarship Council Joint PhD Training Program (D.S.); National Natural Science Foundation of China grant 31630030 (G.B.); Stanford Neuroscience PhD Program training grant 5T32MH020016 and the Post-9/11 GI Bill (S.W.E.); NIH BRAIN Initiative grant 1U01NS090600 (M.J.S. and M.Z.L.); Canadian Institutes of Health Research grant MOP-81142 and Natural Sciences and Engineering Research Council of Canada grant RGPIN-2015-06266 (K.T.); a graduate fellowship from the Natural Sciences and Engineering Research Council of Canada (S.C.); a postdoctoral fellowship from Agence Nationale de la Recherche grant ANR-10-LABX-54 MEMO LIFE (V.V.); La Fondation pour la Recherche Médicale grant Equipes FRM DEQ20140329498 and Agence Nationale de la Recherche grant ALPINS ANR-15-CE19-0011 (S.D.); and additional funding from INSERM, CNRS, and ENS (J.B., B.M., V.V., and S.D.). We also acknowledge the assistance of the Stanford Neuroscience Gene Vector and Virus Core facility, which was supported by NIH grant P30NS069375, and the IBENS Imaging Facility, which was supported by grants NERF 2009-44 and NERF 2011-45 from the Région Ile-de-France; grant DGE 20111123023 from the Fondation pour la Recherche Médicale; a 2011 grant from the Fédération pour la Recherche sur le Cerveau and Rotary International France; and grants ANR-10-LABX-54 MEMO LIFE, ANR-11-IDEX-0001-02 PSL* Research University, and ANR-10-INSB-04-01 France-BioImaging Infrastructure from the Agence Nationale de la Recherche Investissements d’Avenir program.

## Author contributions

M.Z.L. and S.D. conceived the study and designed research. M.C., I.K.D., and L.P. developed the electrical screening platform. M.C., L.P., D.S., and S.W.E. performed library construction, library screening, and indicator characterization in cultured cells. J.B. and S.D. characterized ASAP indicators kinetics at 33°C in cultured CHO cells. S.W.E. and R.Y. performed the acute striatal slice experiments. S.C. characterized indicators in organotypic slices. J.B. designed the viral transgenesis strategies for sparse ASAP expression J.B., B.M. and S.D. performed the acute slice experiments. V.V. and B.M. performed the *in vivo* experiments. V.V. and S.D. analyzed *in vivo* data. S.D. and B.M. conceived and implemented the modified modes of RAMP microscopy. F.S.-P., K.T., G.B., M.J.S., J.D., S.D., and M.Z.L. provided ideas and advice. M.C., V.V., J.B., S.D., and M.Z.L. wrote the paper.

